# NomadFlow: unindexed multi-motif scaffolding with independently mobile rigid groups

**DOI:** 10.64898/2026.10.01.754115

**Authors:** Ivan Permyakov, Elizaveta Geraseva, Andrey Golovin

**Author notes:** Correspondence: Ivan Permyakov, Faculty of Bioengineering and Bioinformatics, Lomonosov Moscow State University, Leninskye gory 1, Building 73, 119234 Moscow, Russia.

## Abstract

Motif scaffolding builds a protein around a prescribed structural motif — an active site, a binding epitope, a metal site — so that the motif is presented in its required geometry. The task is usually posed with the motif pinned down: its position along the sequence is given, and several rigid fragments inherit their mutual pose from a native structure. Many applications supply neither. We present **NomadFlow**, unindexed multi-motif scaffolding on all-atom SE(3) flow matching: the motif enters as a condition, a group projection preserves the internal geometry of each group exactly, and the groups move freely relative to one another until a set time, after which they act only as a condition. We assemble a preliminary benchmark for this setting, varying the granularity of the partition and the structural context around the motif, and obtain unique successes on 19 of 21 tasks. The trained model proposes a diversity of valid mutual poses, and on several tasks the native arrangement is among them. The shape of that solution space, however, varies strongly with the geometry of the condition: across tasks the successful layouts form a wide cloud, collapse onto the native pose, or contract to a connected low-dimensional family — a continuum of poses rather than a set of isolated solutions. Multi-motif methods are therefore better compared by the space of layouts they propose and the successful part of it they retain than by yield or fold diversity alone. Code and weights: https://github.com/ipermyakov/nomadflow.

**Significance statement:** A protein built around several functional fragments is given the shape of each piece but not their relative placement. The strongest methods are told what the motif looks like, but not where in the chain it goes. We adapt one such model so each rigid group moves relative to the others. On a preliminary benchmark for this setting it succeeds on 19 of 21 tasks, and the freedom it uses depends sharply on the motif.

## 1 Introduction

Functional protein design [1] often begins with a small structural motif — the geometry of an active site, a binding epitope [2], a metal coordination site [3], or a set of residues that a larger scaffold must present. **Motif scaffolding** — building such a scaffold around one or more prescribed motifs — is central to the design of enzymes, binders and other proteins with a specified function [4–6]. In recent years generative models, diffusion models above all, have made the task tractable in practice whenever the motif can be supplied as a fixed structural input and embedded into the generated chain by **inpainting** [7], as in RFdiffusion [8] and Chroma [9]. In that formulation both the three-dimensional placement of the motif and its position along the sequence are given in advance; and when several motifs are assembled into one protein, their mutual pose is commonly fixed in advance as well.

In many applications these assumptions are too rigid. Several functional fragments may come from different proteins, and their relative placement in the final structure — **in space** and **along the sequence** alike — is unknown beforehand and, as a rule, does not reduce to a single “correct” pose. **Multi-motif scaffolding** is then not only backbone generation but also a **placement** problem: the model has to find where in the sequence and in space the groups should end up for a consistent scaffold to be buildable around them at all. Inpainting approaches, which pin the motif positions down, scale poorly to this setting; conditional generation without rigid insertion of the motif [10, 11], conversely, does not guarantee that the prescribed geometry appears in the result at all.

Typical cases that call for **unindexed** placement and **several** mobile rigid groups at the same time:

i. *Catalysis plus targeting*. The geometry of an active site is taken from a known enzyme and, separately, a binding motif against a target (a receptor, a surface protein, a partner) from another complex: the internal structure of each fragment is known, but no common template exists in which the two already sit in one chain. Designs of this shape are assembled by hand today — a recent bifunctional degrader carries a target-binding site and a ligase-recruiting motif on one miniprotein, and the placement of the two was settled by trying several grafting positions and loop lengths [12]. Unlike a pocket and an active site serving *the same* substrate [6], here no single productive geometry dictates the mutual pose: different arrangements are admissible as long as both motifs stay accessible and do not clash.
ii. *Two reactions in one carrier*. A single protein is asked to run two different reactions, with the two active sites taken from different enzymes. Such proteins are built today by grafting a second site into a natural enzyme that happens to offer a suitable cavity, and the approach works: two-site constructs improve efficiency, substrate scope and selectivity over a single site [13], and a transaminase paired with an artificial esterase runs a two-step cascade within one polypeptide [14]. Grafting, however, needs a host structure, and the mutual arrangement of the sites is inherited from it. Assembled instead from two prescribed motifs, neither the order along the sequence nor the mutual placement of the groups is given.

Designs of this kind are being built and validated: three non-overlapping RSV epitopes have been scaffolded onto a single domain of under 130 residues, with all three sites confirmed crystallographically [15]. In both scenarios the input supplies neither a correspondence between motif elements and chain positions nor a template for the mutual pose. In the benchmark, by contrast, the groups are taken from a single experimental structure and are therefore compatible at least in the native layout (see Section 3).

Two related but distinct kinds of solution have recently been proposed for multi-motif scaffolding. **MoDiff** [16] trains a diffusion model with a dedicated **Implicit Matching Module**: motifs are encoded by a separate branch, and the matching of motif features to the generated backbone is supervised explicitly through the attention maps, including a motif reconstruction loss. Over the course of diffusion the model implicitly assigns motifs to positions on the scaffold and thereby designs their relative placement, without the positions being specified by hand as in inpainting. **FADiff** [17] takes a different route: motifs enter the generated structure as **floating rigid anchors** that preserve their internal geometry and move as independent rigid bodies, by averaging the SE(3) updates over the residues of a group; the presence of the motif in the result is thereby guaranteed by construction. The order of the motif residues along the sequence is additionally set by an algorithm at sampling time (a short chain, or a TSP over the current structure), so neither method amounts to fully free unindexed placement.

A parallel line of work develops **unindexed** motif scaffolding: the motif is supplied as a condition, through motif tokens for instance, and the correspondence between motif elements and positions in the generated sequence is learned during training (RFdiffusion2, ProteinZen, La-Proteina, RFdiffusion3 and others [18–21]). These approaches remove the need to fix the order of the motif along the chain and currently give the strongest scaffolding on active sites, relative to indexed and inpainting formulations. In the usual formulation, however, the motif is treated as a **single rigid** object rather than as several mobile groups with independent relative motion.

These approaches differ in what the motif is. Inserted into the generated structure, it is present by construction and its geometry comes out exact for free; supplied as a condition, it is usually treated as a single rigid object. We keep the motif a condition, so that generated residues have to recover it, and at the same time make each group an object of generation in its own right: a rigid body whose pose is produced along with the scaffold, and whose internal geometry is exact by construction. We show that this can be done within all-atom SE(3) flow matching: a group projection holds the geometry, while the correspondence of motif elements to positions along the chain is learned implicitly. We build on **ProteinZen** [19], which already conditions on unindexed motif tokens and represents residues as oriented rigid frames — the interface on which a group projection acts directly. For training and for sampling alike we **separate in time** the placement of the motifs from the generation of the scaffold: group motion ends by *t*_lock_ *<* 1, after which the motif is fixed as conditioning and the scaffold is completed to the end.

### Contributions

- **NomadFlow** — unindexed scaffolding of several independently mobile rigid groups, in which the rigidity of each group holds by construction rather than by a penalty;
- a preliminary multi-group benchmark derived from Protpardelle [22], laid out along two axes: how finely the motif is partitioned, and what structural context surrounds it;
- the result that removing the mutual pose leaves the setting solvable, with the cost falling on yield rather than on the diversity of successful folds;
- the observation that successful layouts form sets of very different shape from task to task, and the argument that this is what multi-motif methods should be compared on.

## 2 Problem formulation

### 2.1 Parameterization

Following the ProteinZen parameterization [19], a protein of *N* residues is given by a sequence *s*_1:*N*_ and three local rigid frames per residue: 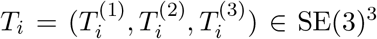. All-atom coordinates are decoded deterministically from the residue type together with these frames. The **generated residues** are the residues of the final chain, whose structure we are building.

A **motif condition** M = {*M*_1_, …, *M*_*m*_} is given, made up of *m* independent rigid groups. Group *M*_*j*_ contains *K*_*j*_ motif residues of known geometry. The model generates **a single chain** 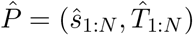 of *N* residues; the motif condition is supplied as conditioning, not as part of the generated structure.

Group *M*_*j*_ is specified by sets of frames 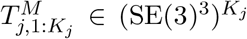. Within a group the relative geometry is fixed, whereas the mutual SE(3) poses of different groups are not given at the input. For *m* = 1 the condition is a single rigid body; for *m >* 1 we obtain multi-group motif scaffolding.

### 2.2 The motif scaffolding task

#### Definition (motif scaffolding)

A motif condition is given as groups *M*_*j*_, *j* = 1, …, *m*, represented by frames 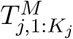. The task is to generate a protein of length *N* — frames 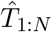 and a sequence *ŝ*_1:*N*_, from which the all-atom structure is decoded — such that the generated residues **reproduce** each group, while the remaining residues form a **scaffold**.

Formally, a successful solution admits injective correspondences *π*_*j*_ : {1, …, *K*_*j*_} → {1, …, *N* } with pairwise disjoint images, and rigid transformations *G*_*j*_ ∈ SE(3), such that 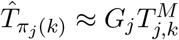 for all *j* and *k*. Here *G*_*j*_ acts identically on all three frames of every residue of the group. A single common *G*_*j*_ for all of its residues expresses the preservation of the internal geometry; the sign ≈ denotes agreement to within a prescribed success criterion. The correspondences *π*_*j*_ are checked after generation, but are neither supplied to the algorithm nor produced as its output. As Figure 1 shows, success means recovering the geometry of the groups through the generated residues, rather than their presence by construction as separate objects of generation.

**Figure 1.**
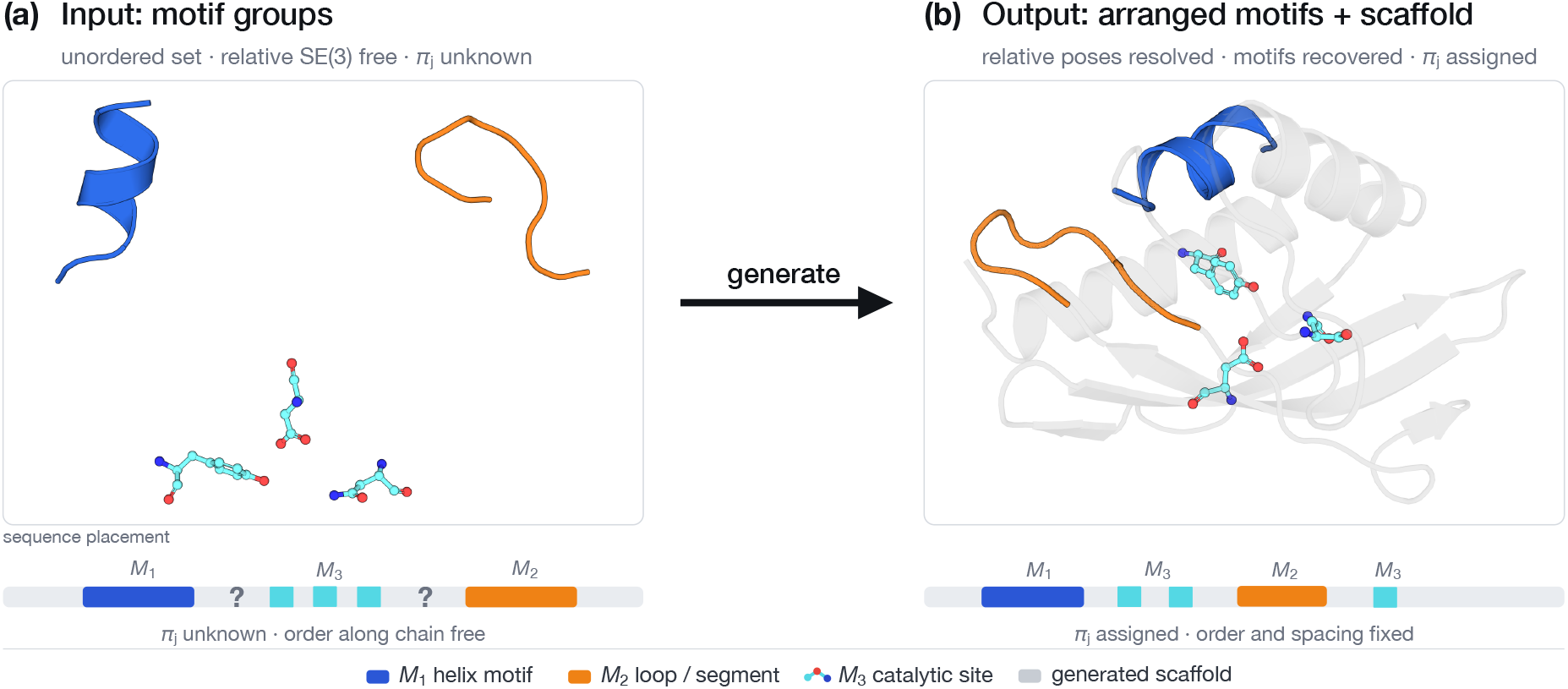
Unindexed multi-group motif scaffolding with several mobile rigid groups *M*_1_, *M*_2_, *M*_3_. *(a)* The input is an unordered set: the internal geometry of each group is given, the relative SE(3) poses are free, and the correspondences *π*_*j*_ are unknown. *(b)* The output is a single chain: the mutual orientation of the groups is consistent with the scaffold, and the groups are reproduced by generated residues. The correspondences *π*_*j*_ are part of the definition of success and are checked after generation; they are not supplied to the algorithm.

## 3 Results

The experimental question of this work is what changes when, for an unchanged set of motif residues, the native mutual pose is removed from the input. To separate the cost of that freedom from any incompatibility of the motifs themselves, we build paired tasks out of groups of a single PDB chain and compare two regimes under one evaluation protocol. **Mobile** is NomadFlow, given the same residues as independently mobile groups; **one-rigid** is the calibration in which the whole set is supplied as one rigid body carrying the native mutual pose (the original ProteinZen). The term *multi-group* describes the setting rather than either regime: the success criterion checks the geometry of each group separately, so the tasks are multi-group in both regimes — one-rigid included, where the groups are fused into a single body at the input. The regimes differ only in whether the mutual pose of the groups is given at the input. The analysis of yield, diversity and mutual layouts that follows rests on this construction.

### 3.1 Method overview: the groups move rigidly, the mutual pose is chosen during generation

The setting (Figure 1) leaves the model free in two respects and fixed in one. The correspondence between motif and generated residues and the mutual pose of the groups are free; the internal geometry of each group is fixed. NomadFlow is arranged so that the first two are resolved over the course of generation, while the third holds by construction rather than by a penalty.

The motif is not inserted into the chain as a block laid out in advance. Each motif residue enters the shared denoiser graph as a separate conditioning token alongside the scaffold tokens, but occupies no position known in advance in the output sequence; in the unindexed regime that position is hidden from the model. The correspondence arises from the spatial coincidence of generated and motif frames and is read off only after generation — there is no separate matching module.

Each group is assigned an auxiliary rigid frame at the center of mass of its motif frames, the **aggregator**. It is not a motif residue but the intrinsic coordinates of the group pose, expressed in the same frame interface the network already uses. The aggregator carries the centroid of the group — the point about which the common rigid update takes place; as a token of the shared graph it passes that point on to the motif tokens through attention.

The rigidity of the groups is enforced by a **projection**. The denoiser produces increments for each motif frame independently; within a group these are averaged into a single SE(3) transformation and applied to all of its frames at once, the aggregator included. The internal geometry of the condition is therefore preserved exactly rather than approximately: there is nothing inside a group that could drift apart, and no intra-group geometric penalty is needed. Different groups meanwhile move independently of one another, and the scaffold is denoised freely.

The training signal for the group pose lives on the aggregator, not on the individual motif residues. The pose of a group has six degrees of freedom; learning it as the independent motion of every motif residue would mean learning something other than the process by which the group actually moves. The aggregator holds the common step of the group and serves as the point through which the gradient returns to the motif tokens whose predictions that step was assembled from.

Finally, the two subproblems — choosing the mutual arrangement of the groups and completing the scaffold — are separated in time by the threshold *t*_lock_ (Figure 2b). Before it the groups are mobile, and the final placement of the motifs is resolved jointly with the coarse topology of the scaffold, so that the pose arrived at is one a scaffold can actually be completed around. After it the groups are fixed and act only as a condition, and the scaffold alone is denoised — the finer adjustments that realize the placement already chosen.

**Figure 2.**
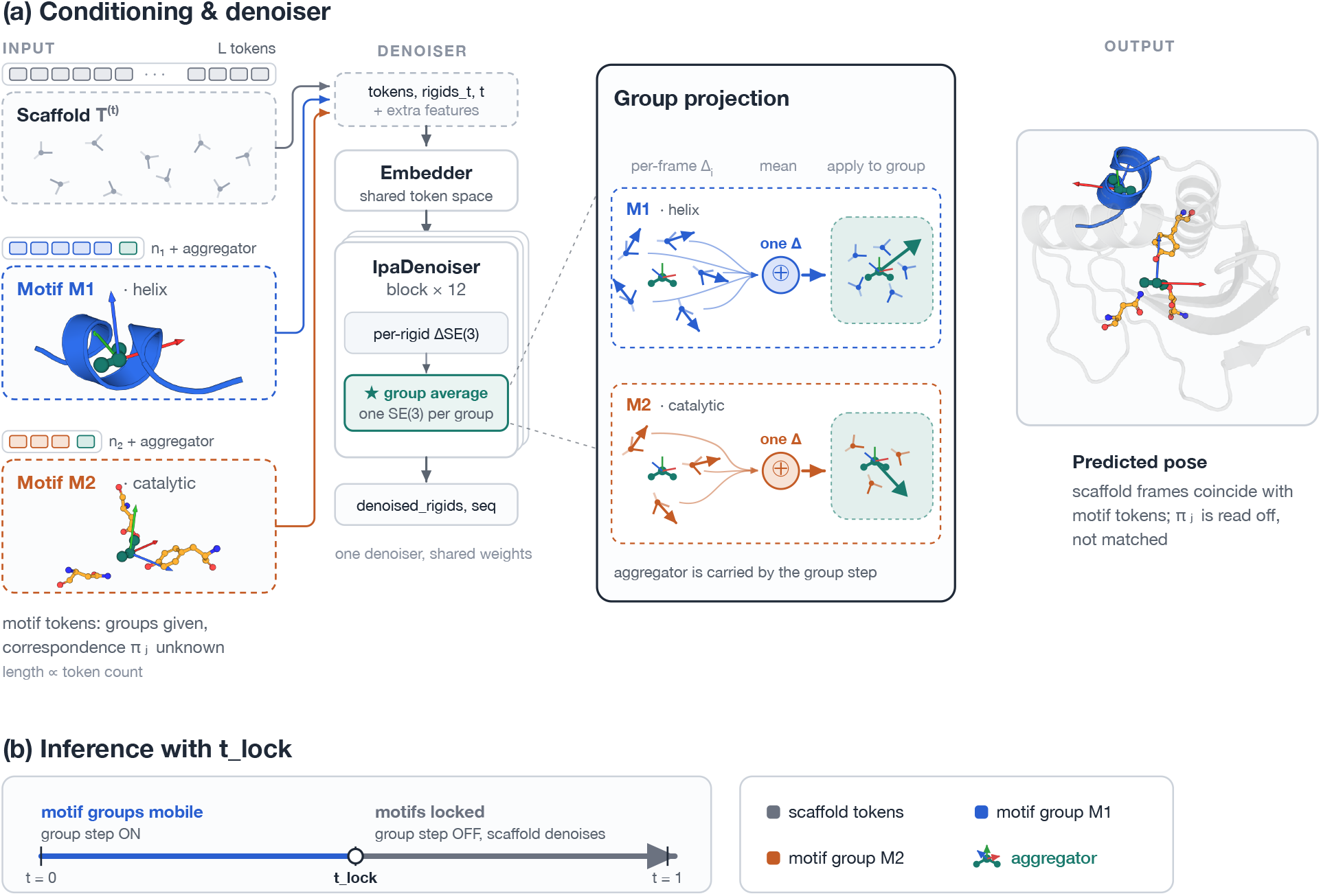
Method overview. *(a)* Motif groups are supplied as unindexed motif tokens (group membership is given, the placement *π*_*j*_ is not) together with the scaffold tokens and one virtual aggregator rigid per group, into a shared denoiser. Per-frame SE(3) increments within each group are averaged into a single rigid step and applied to all frames of the group, the aggregator included: each group moves as a rigid body while the scaffold is denoised freely; the aggregator is carried by that step rather than being its source. The spatial coincidence of generated and motif frames yields the correspondence post hoc, with no explicit matching module. *(b)* Before *t*_lock_ the motif groups are mobile (the group step is on); after *t*_lock_ they are fixed as a condition and only the scaffold is denoised.

The formal account — parameterization, projection, objective and integrator — is given in Section 5.

### 3.2 A benchmark of multi-group tasks

The standard motif scaffolding set [22] specifies the motif as **a single rigid body**: however many fragments it contains, they are immobile relative to one another, and their mutual spatial arrangement is inherited from the native structure. The positions of the fragments along the sequence are not fixed — this is the unindexed setting at *m* = 1. The quantity our setting asks the model to choose for itself — the mutual pose of the groups — is supplied there from the start. The original set does not exercise the *m >* 1 regime — nor do the standardized benchmarks assembled for motif scaffolding since [23] — and we build our tasks from it.

There is a single construction rule. From the same native chain we take the same set of residues as in the original task and declare it to be several **independent** rigid groups. The internal geometry of each group is passed to the model; the mutual pose of the groups is not. The native arrangement remains known to us but not to the model, and does not enter the success criterion. The set of residues is thereby held fixed relative to the original task, and exactly one thing varies — how much information about the mutual arrangement is retained.

The set is organized along two axes, each with a control of its own.

#### Structural segments — the granularity axis

Seven proteins (1PRW, 5IUS, 2KL8, 4JHW, 6VW1, 5YUI, 1BCF); the motif is assembled from continuous fragments of 3 to 52 residues (Figure S1). The set of residues is fixed and the partition into groups varies: 1PRW, 5IUS, 2KL8, 4JHW and 6VW1 run at *m* = 2, 5YUI at *m* = 3, 1BCF at *m* = 4; in this full split each group is exactly one fragment. For 5YUI and 1BCF we add `_2grp` variants — the same set of residues collected into two groups instead of three and four. There, fragments adjacent by contig are fused into one group with a prescribed mutual pose: in 5YUI one pair is fused this way, in 1BCF two. These pairs are the granularity control: within a single protein, with the set of residues unchanged, the only thing that varies is how many mutual poses the model chooses for itself.

#### Active sites — the structural context axis

Four families (1QJG, 5AOU, 5AOU_QUAD, 7K4V; Figure S2). The first group in each is an active site of 3–4 residues **spread along the chain**: in 1QJG, for instance, these are `A38, A14, A99`, three separate residues rather than a fragment. The mutual geometry within the active site is given, so this group alone already binds several fragments into one rigid body.

The zero point of the axis is the original Protpardelle task, in which the motif reduces to a single active site with no structural context (`ext0`); there is one group there, and only one-rigid sampled that task. The reference point for our own moves is `ext10`: the active site plus one continuous segment of ten residues, two groups. Two moves diverge from it, each adding about ten further residues of structural context to the motif, but spending them differently.

**Adding a fragment** (`ext2x10`) leaves the original segment in place and puts a second one of the same length beside it: the residue set of ext10 is contained in ext2x10 in full. There are three groups instead of two, and the model chooses one more mutual pose.

**Lengthening** (`ext20`, and e`xt16` for 1QJG) keeps the number of groups at two but replaces the short segment with a single longer one: no new free poses appear. The long segment is in general taken from a different place in the chain and does not contain the short one; nesting occurs only in 7K4V, where `A61-70` lies inside `A61-80`.

That is how this axis is controlled. Both moves spend a comparable budget of residues, but one goes into the length of a single fragment and the other into the independent mobility of a second one, with `ext10` giving both a common reference point.

| axis | held fixed | varied | tasks | $m$ |
| --- | --- | --- | --- | --- |
| structural segments | residue set within a protein | partition into groups | 9 | 2–4 |
| active sites | active site of the family | length and number of added segments | 12 | 2–3 |

We sample 200 designs per task. The composition of the groups (*F, m*, contig) is given in Tables S1–S2; per-task *S* and *U* are in Tables S3–S4.

This benchmark design is a step toward analyzing the granularity of the partition and the character of the structural context around the active site separately, not a completed factorization. The particular values — the number of groups, the lengths of the additions — were chosen from a sense of what ought to be difficult rather than calibrated in advance; the point of the construction is that the behavior of the model along each axis is measurable on its own. Further versions of the set are naturally built from the results: as shown below, one of the axes does not behave the way its construction assumed.

### 3.3 How often each group is reproduced on its own

Task success is all or nothing: a design counts only if every group comes out right, which leaves open which group was the obstacle. The same data answer a narrower question — how often a single group survives the pipeline, counted on its own.

A design counts for a given group if at least one of its eight ProteinMPNN sequences [24], once folded, reproduces that group to within a heavy-atom RMSD of 2.0 Å and a C*α* RMSD of 1.0 Å of the native geometry — the same per-group threshold the task-level criterion applies. The fold is compared with the geometry that was prescribed, not with the design that was generated, so what is measured is recovery of the motif rather than self-consistency.

Table 1 reports this over the active-site tasks with additions at *m* = 2, where the two groups play clearly different roles: group 1 is the active site, 3 or 4 residues spread along the chain, and group 2 is one continuous segment. Table S2 lists which tasks fall into each row. Every task contributes *N* = 200 designs, so micro-and macro-averaging give the same numbers.

**Table 1.** Fraction of designs in which the given group role is reproduced relative to native (heavy-atom RMSD *<* 2.0 Å, C*α* RMSD *<* 1.0 Å) by at least one of the eight ProteinMPNN sequences. Active-site tasks at two groups, *N* = 200.

| group role | one-rigid | mobile |
| --- | --- | --- |
| active site, 3 residues | 38.4% | 30.0% |
| active site, 4 residues | 40.8% | 32.2% |
| segment, 10 residues | 70.1% | 74.0% |
| segment, 16 residues | 59.0% | 34.5% |
| segment, 20 residues | 87.7% | 66.8% |

The gap between the two roles is wide and survives in both regimes. Under one-rigid the active site is recovered in about 40% of designs against 59–88% for a continuous segment, and letting the groups move lowers the active-site figure further still, to 30.0% and 32.2%. Segments behave less uniformly: mobility is expensive at 16 and 20 residues (59.0% falling to 34.5%, 87.7% to 66.8%) but not at 10, where mobile is marginally ahead (74.0% against 70.1%). The hard part is therefore the active site itself. Holding a handful of chain-distant residues in an exact mutual geometry through folding is difficult in either regime, whatever the model was told about their arrangement.

### 3.4 Both axes are solvable; the cost falls on yield, not on diversity

The first question about the setting is whether it can be solved at all. Under the single criterion described in Section 5, unique success is obtained on 19 of the 21 multi-group tasks: eight of nine on structural segments (nine tasks, two of them `_2grp` variants) and eleven of twelve on active sites. Two are missed: `4JHW`, where mobile produced no successful design at all (*S* = 0) while one-rigid produced some (*U* = 3), and `7K4V_ext2x10`, where both regimes give zero (*S* = 0, *U* = 0).

Successful designs from both axes are shown in Figure 3.

**Figure 3.**
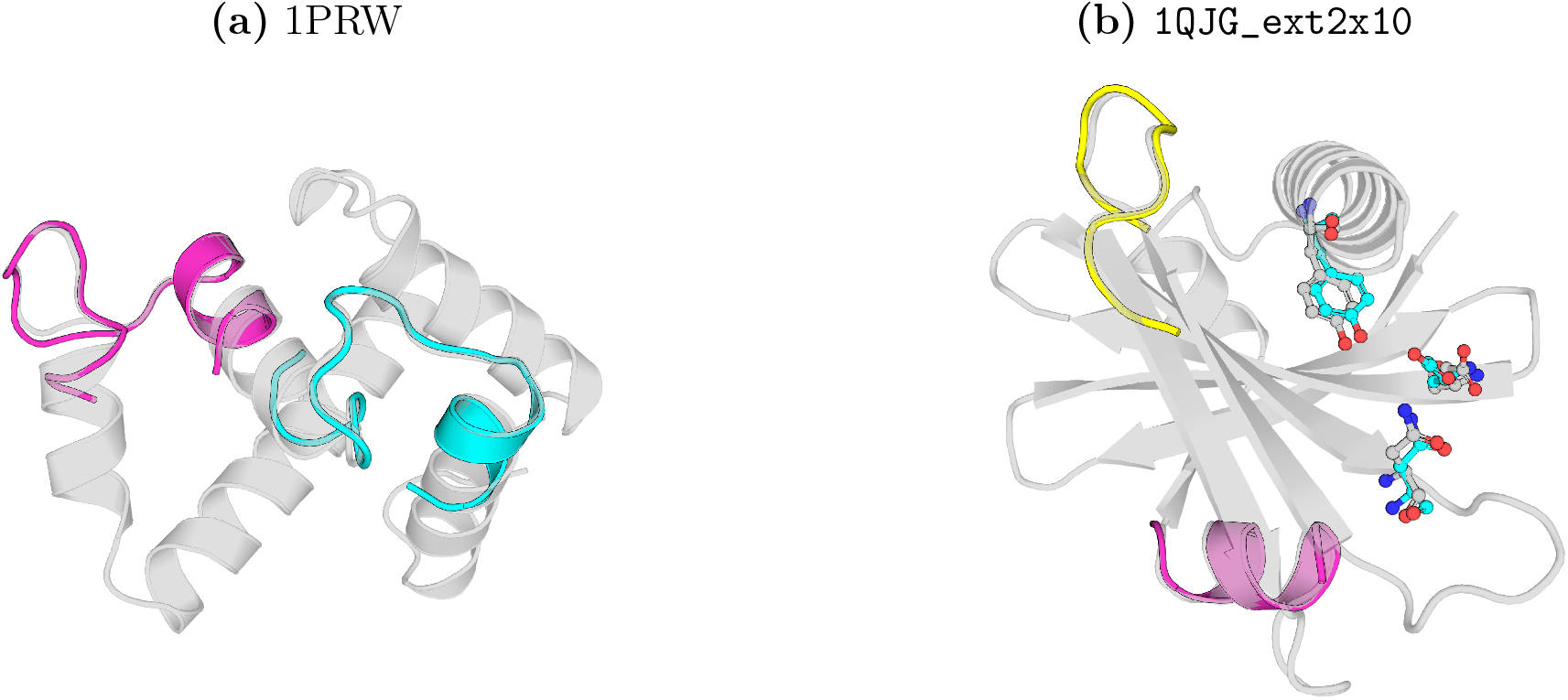
Successful NomadFlow designs after ProteinMPNN and ESMFold. The scaffold is gray; the motif groups are colored; the crystal geometry is superimposed on the motif sites of the folded structure (heavy-atom alignment). *(a)* 1PRW: two segments of 20 residues each (A16–35 and A52–71). *(b)* `1QJG_ext2x10`: the active site {A38, A14, A99} as sticks, plus two segments A22–31 and A103–112.

Each mobile task is paired with one-rigid, the unindexed setting at *m* = 1, where the demand on the model is strictly weaker: the same benchmark, the same criterion, the same set of residues, but the mutual arrangement of the groups may be fixed to native. It sets the scale for that easier specification and allows a descriptive comparison with mobile on the same motifs, partitioned into independently mobile groups.

On the structural axis (Figure 4) the tasks fall into two distinct groups by the observed scale of unique. On 1PRW, 6VW1, 5YUI and 1BCF at least one specification gives between 27 and 112 unique successes, and on all four proteins the full split exceeds one-rigid. On 5IUS, 2KL8 and 4JHW neither value exceeds three clusters: 2 →1, 2 →2 and 3 →0 respectively. These tasks enter the aggregate figures, but at values this small they say almost nothing about the direction of the difference in diversity.

**Figure 4.**
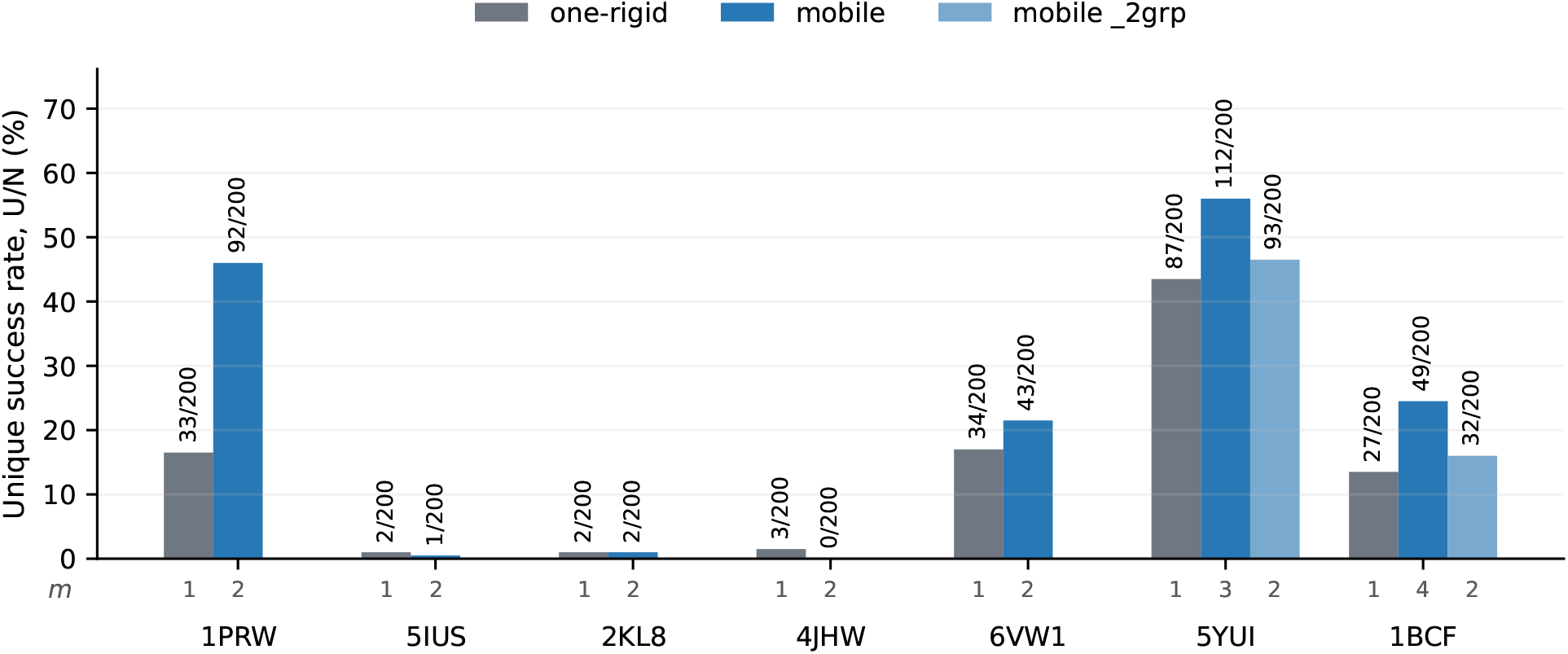
Unique success rate *U/*200 after ProteinMPNN and ESMFold on seven proteins, with the same set of residues within each protein. Gray is one-rigid. Blue and light blue are two partitions of the mobile regime: the full split, where each group is exactly one fragment, and the `_2grp` variant, where adjacent fragments are fused into two groups (5YUI and 1BCF only, not separate proteins averaged). The figures beneath the bars are the number of groups *m*; for one-rigid it is 1 throughout. A design is successful if at least one PMPNN sequence reproduces every group relative to native (heavy-atom RMSD *<* 2.0 Å, C*α* RMSD *<* 1.0 Å). Unique counts Foldseek clusters (TM 0.5) among these successful design backbones. *N* = 200; bar labels are *U/*200.

The `_2grp` variants give a further comparison of granularity at an unchanged set of residues. Partial grouping falls between one-rigid and the full split: 87 →93 →112 on 5YUI and 27 →32 →49 on 1BCF. The informative comparisons on the structural axis therefore show no impoverishment of unique when the groups move separately; on the contrary, in the cases observed, more free mutual poses correspond to more unique successes.

On the active-site axis (Figure 5) `ext0` in each series gives the active site with no structural context. There is only one group there, so no paired comparison with mobile exists; a non-zero unique in all four series shows that the active sites themselves admit successful designs.

**Figure 5.**
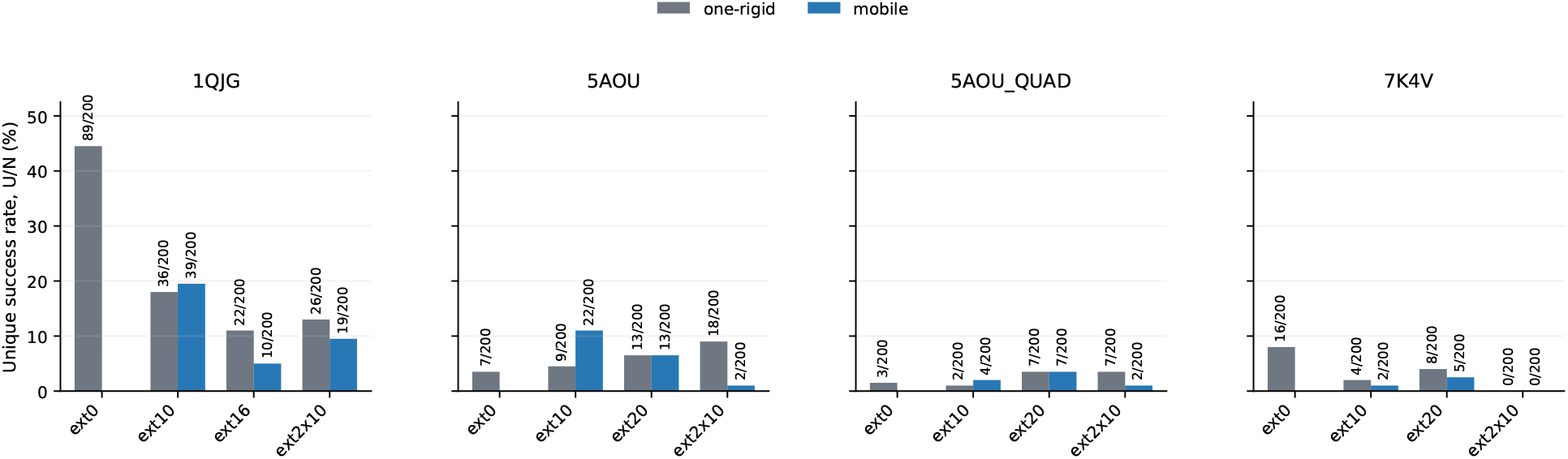
Unique success rate *U/*200 for four active sites under different variants of structural context; the protocol is the same as in Figure 4. Tasks with no added segments (`ext0`) were sampled by one-rigid only: this shows how much added segments in known positions complicate the task even with the native pose fixed. On the tasks with additions the bars are paired: one-rigid against mobile on the same contigs. For 1QJG the long continuous addition contains 16 residues, and 20 for the other series. *N* = 200.

On the tasks with context the direction of the differences in unique is not uniform. Mobile gives more clusters only in the three `ext10` variants, and a pronounced increase appears only on `5AOU_ext10` (9 →22); the differences on `1QJG_ext10` (36 →39) and `5AOU_QUAD_ext10` (2→ 4) are small. This axis therefore shows no general advantage for either specification in unique. The dependence on the type of structural context is taken up below.

To reduce each pair to a single number per task, we take the ratio of the mobile value to the one-rigid value on the same set of residues — separately for yield *S* and for unique *U* . Of nineteen such pairs, eighteen ratios are finite; `7K4V_ext2x10` gives 0*/*0 and is excluded from the comparison. The median is 0.57 for *S* and 1.00 for *U* . On unique, mobile wins on seven comparable tasks, loses on eight and draws on three. The gap to one-rigid is therefore more pronounced in the frequency of success, while there is no systematic direction in the diversity of successful folds.

Foldseek [25] distinguishes global scaffold folds but says nothing about the mutual poses of the motif groups. The freedom given to the model therefore has to be measured separately.

### 3.5 Successful layouts are diverse and include the native one

For each motif group we define a local frame from its backbone atoms. For a pair of groups *i, j* we take the relative pose 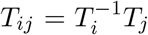, which does not depend on how the structure is placed in space. The set of all *T*_*ij*_ over pairs of groups is the **layout** of a design: precisely what the setting leaves the model to choose. The distance between two layouts is the norm of the difference in *t*_*ij*_, in angstroms, together with the geodesic distance between the *R*_*ij*_ in SO(3), in degrees; pairs of groups are combined as a root mean square, with equal weight and irrespective of group size.

On 1PRW, 6VW1, 5YUI and 1BCF the successful layouts occupy a wide range of poses — tens of angstroms and well past 90^°^ (Figure 6). Of the eight structural tasks with *S* ≥1, four have at least one success within 2 Å and 15^°^ of native: 1PRW, 2KL8, 6VW1 and 5IUS. The native layout is visited, but it is not the only one found.

**Figure 6.**
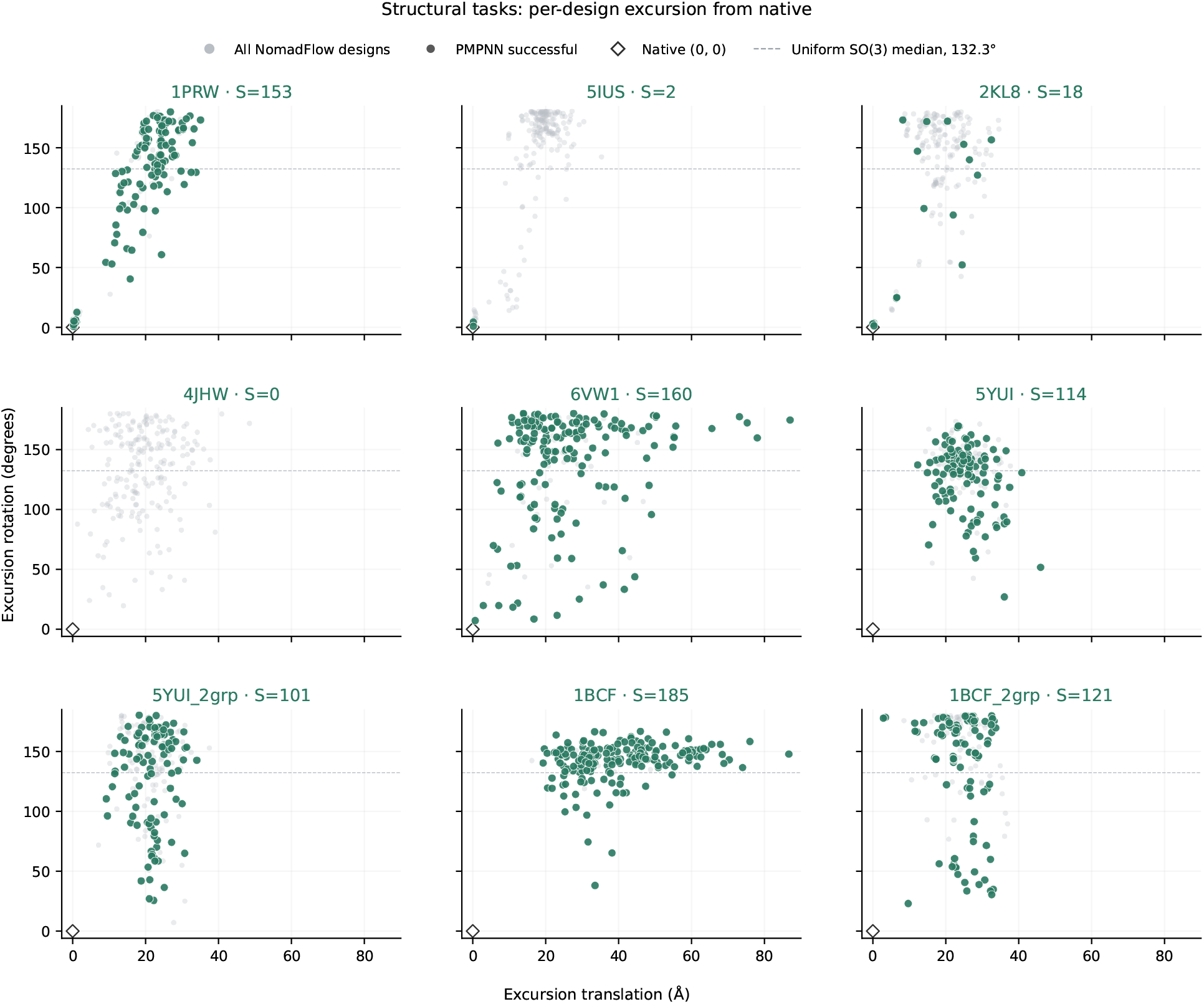
Excursion of the layout from native in translation (Å) and rotation (degrees) across the nine structural tasks. Gray is all 200 NomadFlow designs, color marks the successes after PMPNN+ESMFold, and the diamond is native. *S* in each panel title is the number of successful designs. For *m >* 2 both coordinates are root mean squares over pairs of groups.

On 5IUS the proposals themselves are varied and far from native. Only two designs succeed, both close to the crystal pose (median 0.23 Å / 2.6^°^). The model proposes alternative poses, but only the native one survives PMPNN+ESMFold.

The excursion from native is only one projection of the set of layouts. A second is the pairwise distance between the designs themselves, which shows how far they diverge from one another independently of the crystal: the joint distribution of both coordinates over the structural tasks is given in Figure S3. A summary of the medians of both quantities across all tasks at once, with the step from all designs to the successful ones, is given in Figure S4.

### 3.6 The freedom in pose can contract to a low-dimensional family

With the long continuous segment, on `5AOU_ext20` and `5AOU_QUAD_ext20`, the successful layouts occupy a narrow locus far from native: tens of angstroms and more than a right angle, with 143 and 77 successes. Translation and rotation are coupled; this is not a collapse onto the crystal pose as on 5IUS. Such a set reflects the successes observed for this model under the PMPNN+ESMFold filter, not proof that the task is one-dimensional (Figure 7).

**Figure 7.**
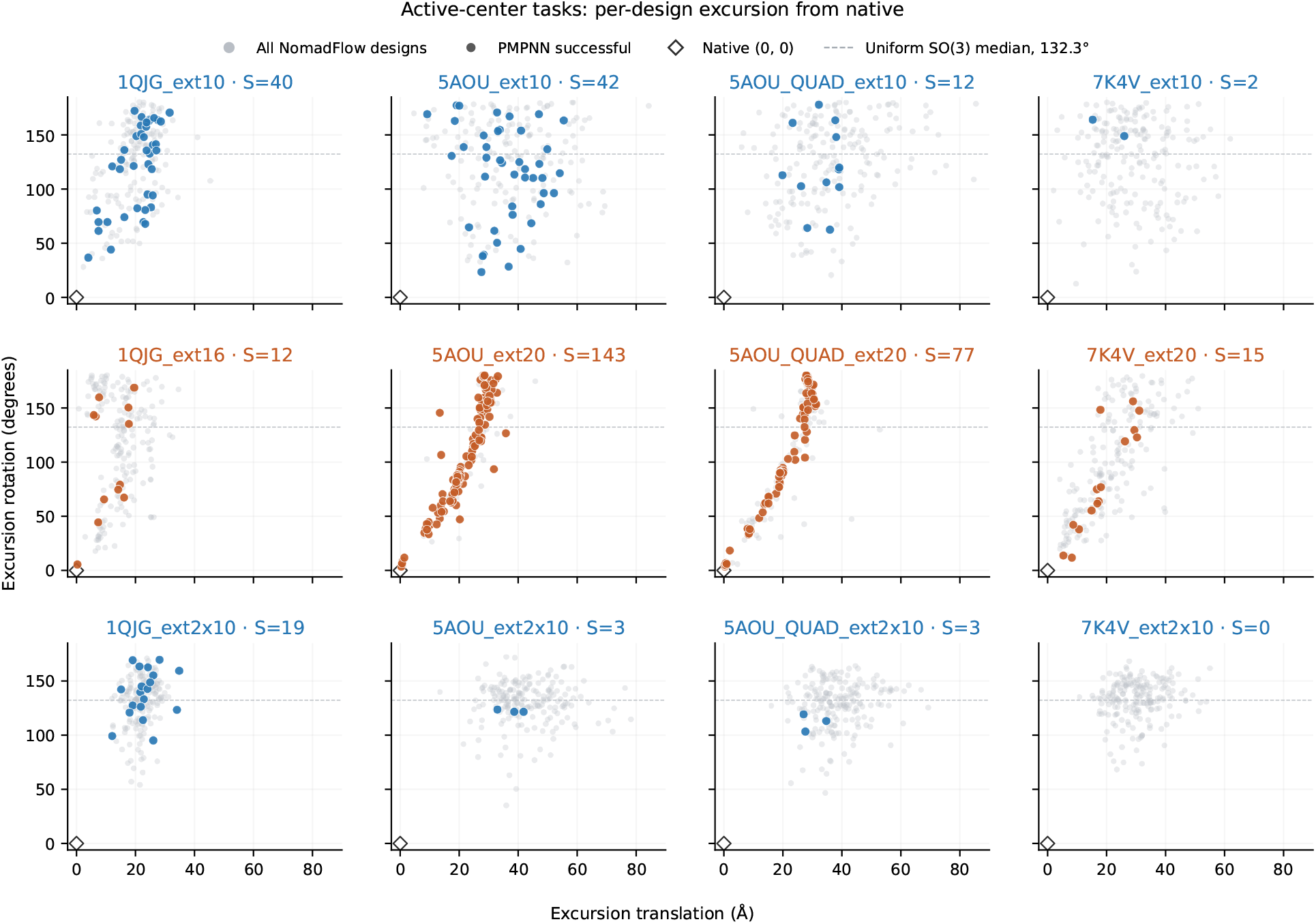
Excursion of the layout from native in translation (Å) and rotation (degrees) for the twelve active-site tasks. Rows are the addition categories `ext10`, the long continuous segment (`ext16` for 1QJG, `ext20` for the rest) and `ext2x10`; columns are the families 1QJG, 5AOU, 5AOU_QUAD and 7K4V. Gray is all 200 NomadFlow designs, color marks the successes after PMPNN+ESMFold, and the diamond is native. *S* in each panel title is the number of successful designs.

On the same proteins the short `ext10` leaves a wide set of successful layouts, while `ext2x10` gives varied proposals with almost no successes. On `1QJG_ext10, 1QJG_ext16` and `7K4V_ext20` the successes also lie far from native in a similar direction, and something of the same outline can be made out — but far less sharply: the cloud stays diffuse, and we do not read a two-dimensional shape into it. Pairwise distances between layouts on this axis are given in Figure S5.

Two-dimensional classical MDS of the same `5AOU_ext20` and `5AOU_QUAD_ext20` successes yields a circle (Kruskal stress [26] 0.12 and 0.15; the two coordinates retain 85% and 82% of the positive spectrum) (Figure 8). For the remaining tasks on the same axis the stress is high or the points are few: we do not read the same circle out of those embeddings. The embedding for the structural axis is given in Figure S6.

**Figure 8.**
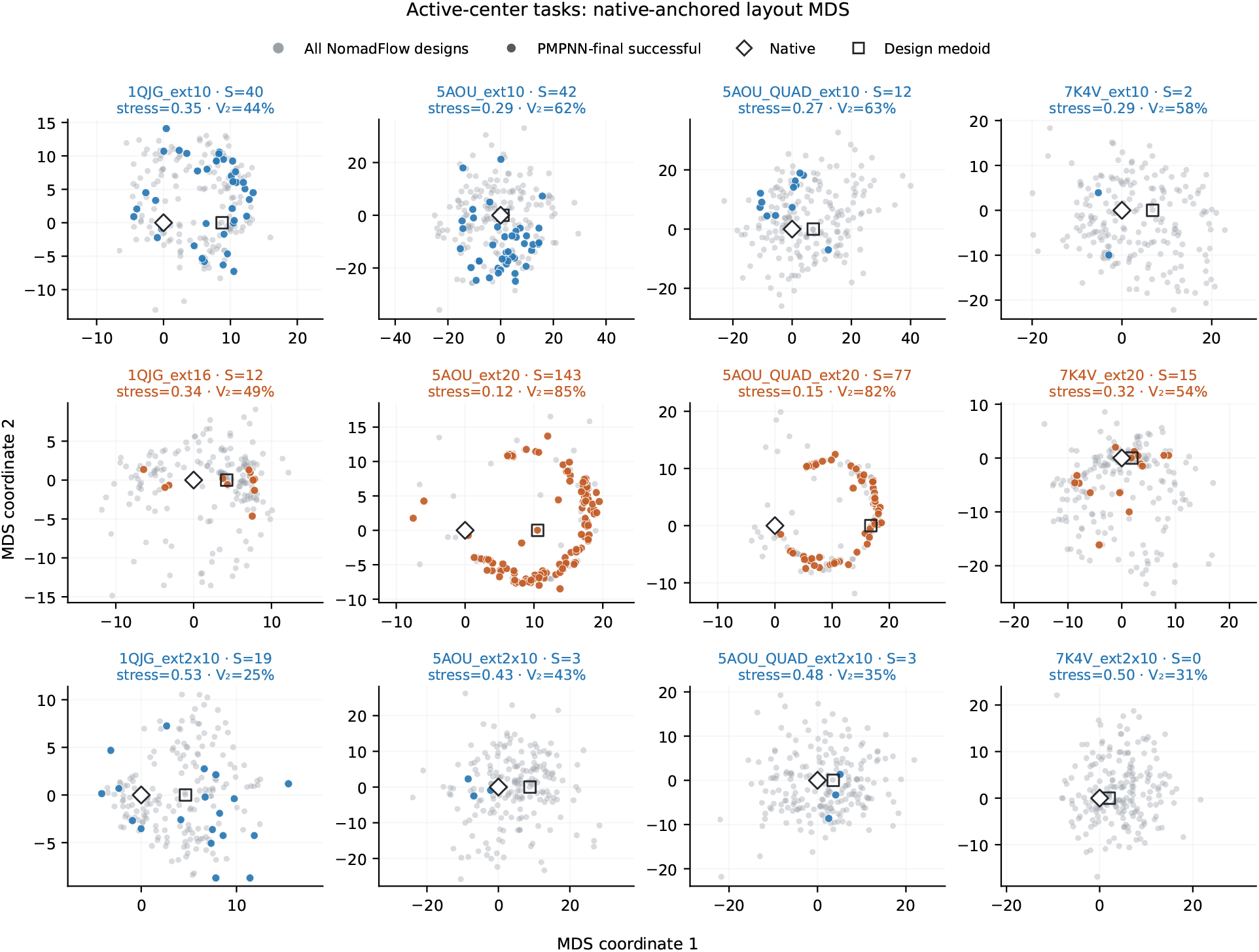
Native-anchored classical MDS of the same twelve tasks. Rows and columns are arranged as in Figure 7, and the point symbols follow it; the square is the medoid. Panel titles give the Kruskal stress and the fraction of the positive spectrum retained by the two coordinates *V*_2_. We do not interpret the shape of panels with high stress.

In three dimensions the same `5AOU_ext20` successes line up along a one-dimensional trajectory of the second fragment across the surface of the protein (Figure 9).

**Figure 9.**
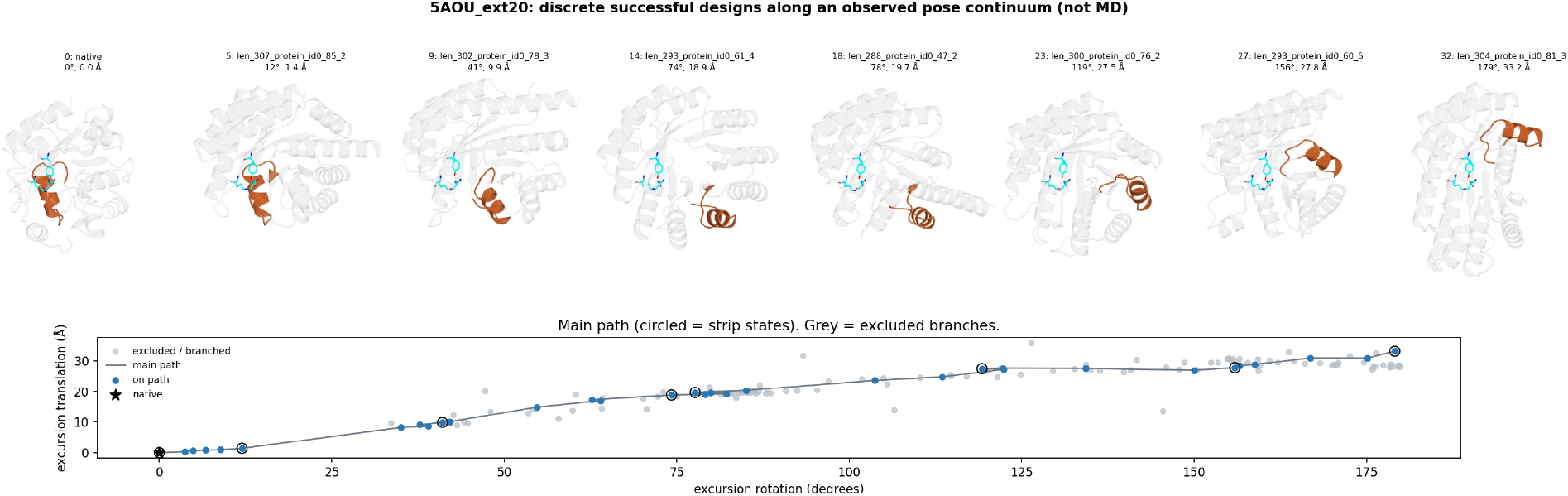
Representative successful `5AOU_ext20` designs. The ordering is a walk through neighboring layouts in the space of excursions from native (the same axes as in Figure 7), from the crystal pose to the most distant success. Laid out this way, the different poses form a one-dimensional trajectory of the second fragment. The active site is superimposed. These are discrete structures, not molecular dynamics.

All the layouts above are measured on the generated designs, whereas success is determined after ProteinMPNN and ESMFold. The median design →fold error among successful designs shows that the layouts are largely retained through folding (Figure S7).

## 4 Discussion

The main result of this work is that unindexed motif scaffolding with several independently mobile rigid groups remains a computationally solvable task. Although the model has to find the correspondence of motif elements to positions along the chain, choose the mutual pose of the groups and build a scaffold around them all at once, unique successes are obtained on 19 of the 21 multi-group tasks. Removing the mutual arrangement from the input therefore does not push the setting into a practically unreachable regime: its cost is finite and can be measured at a fixed generation budget.

That cost shows up first of all in the probability of landing on a successful design. Relative to the one-rigid calibration the median ratio is 0.57 in yield *S* and 1.00 in unique *U* . The freedom in mutual pose therefore does not cut the successful solutions down to a single preferred fold: it makes motifs and scaffold agree less often, while the diversity of successful layouts is on average preserved. The strong heterogeneity between tasks matters here: the cost depends not only on the number of motif residues, but also on how they are divided into independently moving groups.

Across tasks the proposed and successful layouts occupy sets of different dimension and shape (Figures 6–9, 8): a wide cloud, a collapse onto native, a connected low-dimensional family far from the crystal.

Models are therefore more informatively compared not by the diversity of folds alone, but by which subspace of mutual poses a model proposes and which subspace of successful poses it retains out of it.

Under such a comparison, on each task the designs of different methods are clouds of points in one and the same space of layouts, and the measure of coverage is the fraction of the volume of the common atlas of successes that each model covers.

The composition of the benchmark follows from this. It needs tasks of both kinds: some in which the proposed cloud of layouts is wide but few of its poses fold, and some in which the successes are confined to a narrow region. The first kind asks whether a method can locate the rare foldable poses inside a large space of candidates. The second asks what no single model can answer on its own: whether that narrow region is all the task admits, or merely all that one model ever proposes. If several methods converge on the same region, the constraint belongs to the task; if each settles in a different one, what was being measured was the model. A benchmark built from one kind alone loses this distinction and falls back on yield — where almost nothing folds every method looks weak, and where a small region folds readily every method looks strong.

The measure itself is formulated here but not applied across several models. We draw no direct comparison with MoDiff or FADiff. Both contribute the idea, and we position our work against them above; but neither is distributed in runnable form, code and trained weights not having been released together, so neither could be run on our benchmark. A numerical comparison by coverage of the space of layouts therefore remains the nearest continuation of this work, as soon as the methods become available. So do ablations of the individual choices in the method — first of all the value of *t*_lock_, which sets the boundary between placing the groups and completing the scaffold.

The tasks are assembled from a single PDB entry and guarantee at least one compatible layout; they do not test the assembly of motifs drawn from different proteins, and computational success after PMPNN and ESMFold does not imply expression or function. Taken together, unindexed placement of several groups remains feasible, and its cost falls above all on yield. Methods should be compared, and benchmarks assembled, by the geometry of the proposed and successful layouts rather than by a single figure for yield or for the diversity of folds.

## 5 Materials and methods

### 5.1 Method

NomadFlow generates a sequence and an all-atom structure from an unordered set of rigid motif groups, with no mutual poses given at the input. It is conditional flow matching [27] on the ProteinZen backbone [19], which continues a line of SE(3) generative models for protein backbones running from diffusion [28] to flow matching [29, 30] and on to motif scaffolding [31]. Motif residues are supplied as separate conditioning tokens, the internal geometry of a group is preserved by a projection, and the mutual poses are fixed by *t*_lock_ (Figure 2).

#### 5.1.1 The generative model

The parameterization uses three oriented rigid frames per residue. For compactness the equations below are written for a single component frame 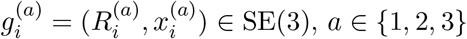, and apply to each of the three frames; the index *a* is omitted from here on. Atoms are decoded deterministically from them together with the predicted amino-acid type, so structure and sequence are predicted jointly, as in other joint sequence–structure generative models [32, 33]. From the noised state at time *t* ∈ [0, 1] the denoiser predicts the clean frame states 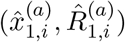.

##### Scaffold

For scaffold residues the path from prior to data is straight in the standard parameterization. Prior: *x*_0,*i*_ ∼ N(0, *I*), with *R*_0,*i*_ on SO(3);

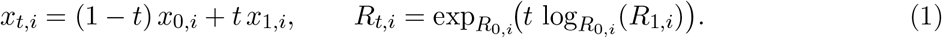

The reference rotation velocity is 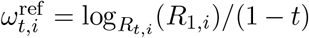, and 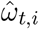 is obtained the same way from 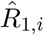. Per-residue FM on the scaffold:

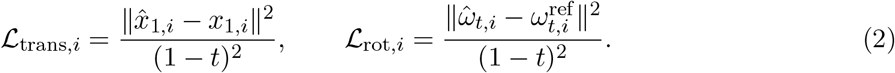

The (1 − *t*)^−2^ scale is a reparameterization of the denoising error on clean data. The sequence loss and a pairwise geometric term, FAFE, enter the same objective as separate terms (the full L is given below). FAFE is inherited from ProteinZen: for every pair of frames it compares the predicted relative transform 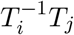 with the ground-truth one, penalizing the deviation in translation, clamped at 20 Å, together with the geodesic deviation of the relative rotation in SO(3). Being built from relative transforms, it is invariant to a global rigid motion of the structure. Details of the architecture and of the integrator are given by Li et al. [19].

#### 5.1.2 The condition: motif tokens and the partition into groups

The motif is not inserted into the generated chain as a fixed block. Each motif residue is supplied as a separate **conditioning token** (motif token): it takes part in the shared denoiser graph alongside the scaffold tokens, but occupies no position fixed in advance in the output sequence. When motif positions are supplied, the model is given *π*_*j*_ directly; when they are withheld — the unindexed case, and the one used throughout this work — the correspondence between motif and generated residues has to emerge in the course of generation.

The partition M= {*M*_1_, …, *M*_*m*_} is given at the input: the tokens of one motif *M*_*j*_ share a common **rigid group motion**, while different motifs may move independently. The condition thus carries the internal geometry of each fragment but imposes neither an order along the chain nor a mutual pose of the groups.

For each group we introduce a **group aggregator**, an auxiliary frame at the center of mass of the motif frames. It is not a motif residue: it holds the intrinsic coordinates of the group pose in the same frame interface the network uses. The aggregator carries the centroid of the group — the point about which the common rigid update takes place; as a token of the shared graph it passes that point to the motif tokens through attention. After the group projection it holds the common SE(3) update; the FM loss is computed on it, and through the projection the gradients reach the motif tokens (see below).

Scaffold tokens and motif tokens are processed by one and the same denoiser: they are distinguished by their role in the condition, by the noising masks and by group membership, not by a separate branch of the network. Membership of a motif group is encoded at the token/frame level and as a pair type (within a group / between groups / motif–scaffold); the aggregator flag is a separate frame feature.

#### 5.1.3 Rigid motion of a group

At the end of the IPA block [34] the denoiser predicts frame updates individually. For a motif group they cannot be applied independently, or the prescribed internal geometry would drift apart. The per-frame increments of the members of a group are therefore first averaged, in the **global** frame, into one common SE(3) transformation, and it is this common update that is applied to all frames of the group — the aggregator included.

Let R_*g*_ be the motif frames of group *g* (the aggregator excluded). Write 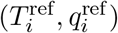 and 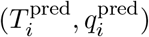 for the states before and after the per-frame update. For each *i* ∈ R_*g*_ we take the rotational increment in the global frame 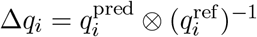 and average the quaternions on *S*^3^ with sign alignment: 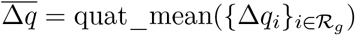. Let 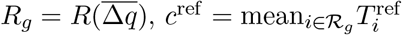 and 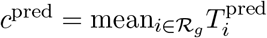. Then for every frame *j* of the group, the aggregator included,

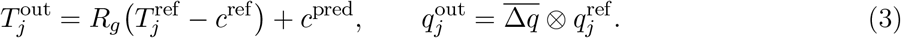

For the aggregator itself, which lies at the center of mass 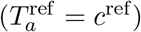, the translation reduces to 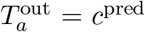: what remains in it is exactly the common shift and rotation of the group, with no intra-group displacement relative to the centroid.

Within a group the geometry is preserved exactly; between groups the transformations are independent. The same projection is used in training and in sampling alike: the source of the motion is the averaged updates of the motif frames, and the aggregator **stores** that common update and serves as the point of supervision (see below) rather than as a separate generator of the group’s motion.

#### 5.1.4 Training

For mobile motif groups the aim of generation is to carry the orientation and position of a group from the prior to the target pose of the conditioning: *g*_0_ ∼ *p*_0_ → *g*_1_, where *g* ∈ SE(3) is the rigid pose of the group. Flow matching requires **training trajectories** for this, and we want them to be linear in the standard SE(3) parameterization. Linearity is taken not along the individual motif frames but in a coordinate system attached to the **center of mass** of the group: the path is a single rigid motion *g*_*t*_ of the group as a whole, from a random *g*_0_ to the target *g*_1_, and the motif residues are carried rigidly along with the group. These are the paths we use in training, and, consistently, in sampling up to *t*_lock_.

Learning this placement of groups and the completion of the scaffold at the same time, along one global axis *t*, mixes two different subproblems. We separate them in time by a threshold *t*_lock_ ∈ (0, 1). The local time of the group motion is

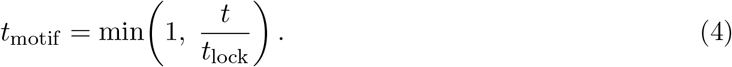

While *t < t*_lock_, each group follows the path *g*_*t*_ described above in local time *t*_motif_, whereas the scaffold follows the ordinary *t*. At *t≥ t*_lock_ the motif frames and the aggregators are fixed in the target state and are from then on only a condition, no longer an object of the transformations; the scaffold goes on being denoised until *t* = 1. The early phase is thus responsible for finding the mutual poses of the groups and for a coarse agreement between motifs and scaffold that the later completion can build on; the late phase is responsible for bringing the scaffold into exact agreement with a motif arrangement that is already fixed.

##### Loss for mobile motif groups

For the scaffold the basic objective is already given above: the denoiser predicts clean frames, the path is straight, and per-residue translation/rotation FM carries the (1 ™ *t*)^−2^ scale. Per-residue FM cannot simply be carried over to the frames of motifs that move as a group, however. For the pose of a group we also want a straight path — but in the coordinates of *g*_*t*_, not along each motif frame separately.

Under rigid motion of a group the individual paths of the residues are no longer straight: they are induced by a common SE(3) element acting about the center of mass. In the ambient space of the frames of a group, SE(3) ^|*g*|^, the admissible configurations lie on a **submanifold** of rigid poses of dimension 6 rather than 6·|*g*| ; the probability measure on the ambient space is singular. A plain independent per-residue FM, written out as it is for the scaffold, therefore matches neither our noising and sampling process for a group nor the geometry of the space on which the data are concentrated.

There are two ways to bring the objective into line with this process. The first is to stay in ambient coordinates and decompose the group motion explicitly into an angular velocity *ω*(*t*) and a velocity of the center of mass ċ(*t*), accounting for the path-correct velocity at every motif frame. The second is to move to the **intrinsic coordinates** of the group: to the single pose *g* = (*R, c*) ∈ SE(3), on which the path is already linear, and to apply ordinary denoising FM there.

We take the second route: it follows directly from the choice of training trajectories *g*_*t*_. The aggregator represents *g* in the frame interface of the network. The standard FM loss is placed on it, and the real motif frames are excluded from the FM loss. The weight *w*_agg_ is assigned **per group** — a constant, not ∝ |*g*| . Gradients from this loss pass back through the application of the common update and through the averaging of the increments (see above), supervising the predictions of the motif frames out of which the update was assembled. The rigidity of the members of a group is fixed by construction (the group projection), so the dynamics stay on the intended submanifold. An intra-group geometric penalty is redundant: by construction there is no drift within a group.

The pairwise geometry (FAFE) is split into three roles:

- **base FAFE** — as in ProteinZen, mainly scaffold and motif–scaffold;
- **aggregator FAFE** — pairs involving aggregators (the pose of a group relative to the remaining frames);
- **intergroup FAFE** — for *m >* 1, pairs of motif frames with group_*i*_ ≠ group_*j*_ (the relative arrangement of the groups; pairs within one group are not penalized).

Formally, the set of frames I_flow_ used for translation/rotation FM contains the scaffold and the aggregators and excludes the real motif frames:

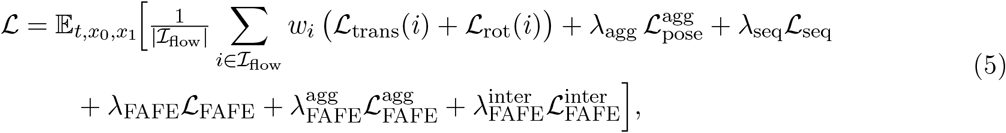

where *w*_*i*_ = *w*_agg_on the aggregators and 1 on the scaffold, and 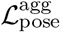is the FM on the aggregator pose. The scaffold error is scaled as (1 − *t*)^−2^. For the aggregator and for 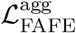 and 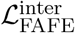 the same scale is taken in the time of the group path, (1 ™ *t*_motif_)^−2^, since *x*_*t*_ of a group is built in *t*_motif_ and not in global *t*. These group terms are also multiplied by 1*/t*_lock_: placement is alive only up to the lock, and with *t* ∼*U* [0, 1] their contribution would otherwise be drowned out by the scaffold. This aligns the two schedules. Numerical values of the weights are given in Table 3 (Appendix B).

Learning the independent motion of several groups requires training examples with several motifs on one chain, and these cannot be assembled by an arbitrary crop. Two naive orderings break a multi-group training example: cropping the chain first and choosing the motif afterwards may leave the fragment without room for the groups, while choosing the motif on the full protein and then cropping independently lets the crop cut through a group that has already been chosen. On the full protein, therefore, farthest-point sampling over C*α* divides the chain into spatial cells, a motif candidate is chosen in each of them in advance, and a motif enters an example only together with its whole cell. The benchmark partitions are specified separately at evaluation time and do not coincide with this construction. The algorithm is given in Appendix A.

The final NomadFlow checkpoint is a finetune of pretrained ProteinZen rather than training the whole network from scratch: the remaining weights are taken from the pretrained checkpoint, and only the new embeddings are initialized afresh. Finetuning proceeds in two stages of increasing crop size, with the task mixture and *t*_lock_ unchanged: only the length of an example grows, so the second stage sees more groups per chain and longer scaffolds. The total length of finetuning is a small fraction of pretraining, so group motion comes to the pretrained network comparatively cheaply. The budget, learning rate and schedule are given in Appendix B.

#### 5.1.5 Sampling

Sampling is the reverse integration of the same flow along the global *t*, with a stochastic integrator. At every step the motif tokens and the aggregators enter the denoiser as a condition. Before *t*_lock_ the groups are mobile: once the frames have been updated, the same projection as in training assembles the common step and applies it to the whole group, the aggregator included, and the group step runs in *t*_motif_ . After the lock the poses are fixed and only the scaffold is denoised. How the group step is reconciled with the two times is set out in Appendix C.

In all runs reported here *t*_lock_ = 0.5, the same value as in finetuning, so half the trajectory goes to placing the groups and half to completing the scaffold. The stochastic noise on motif groups obeys the same rigidity: per-rigid churn is switched off on them and replaced by a single common SE(3) noise per group, since independent noise per residue would destroy the prescribed internal geometry. The integration grid and the churn constants are given in Appendix C.

The output is a single chain. The network does not predict the correspondences *π*_*j*_ explicitly: after generation the motif and scaffold residues end up close together in space, and the correspondence is read off as the nearest pair.

### 5.2 Evaluation protocol

Both regimes are evaluated under one protocol, taken from the ProteinZen evaluation on the Protpardelle benchmark [19, 22] and modified in the two respects noted below; validating designs by sequence redesign followed by structure prediction is standard practice in motif-scaffolding benchmarks [8]. **One-rigid** is the original ProteinZen, given the same set of residues as a single rigid group with the native packing retained. **Mobile** is NomadFlow: the same motifs, partitioned into independently mobile groups. Motif groups are annotated in both campaigns and the same success criterion is applied to both: what differs is the freedom during sampling, not the definition of a successful fold.

We sample 200 designs per task. The chain length *N* is not fixed: it is sampled independently for each design from a range set per task (Tables S1–S2, column *L*). The ranges are inherited from the original Protpardelle tasks and shifted by the length of the added segments, so that within a one-rigid / mobile pair the length is distributed identically and does not interfere with the comparison. **The main protocol of this paper is ProteinMPNN + ESMFold** [24, 35]. One-shot all-atom self-consistency is deferred to the supplement (Figures S8–S9) and is never combined with PMPNN into a single number. One-shot serves as a self-consistency control, not as the main protocol.

For each design we generate eight ProteinMPNN sequences. A design is successful if at least one of them reproduces every motif group relative to native (heavy-atom RMSD *<* 2.0 Å, C*α* RMSD *<* 1.0 Å) — the same criterion as in the caption of Figure 4. Each group is superimposed separately, so the mutual arrangement of the groups does not enter the binary success: closeness of the inter-group arrangement to native is a permitted degree of freedom, not a requirement. The pooled (whole-motif, global) RMSD is computed as a diagnostic and does not enter the all-groups criterion.

The criterion departs in two ways from the one it is taken from. It is applied to each group separately and conjoined over groups, since the setting supplies several groups rather than a single motif. And the global RMSD term is dropped: what is asked here is whether the prescribed geometry is reconstructed in the predicted fold, and agreement between the predicted structure and the generated design over the whole chain is not required. Success rates are therefore not directly comparable with published numbers obtained under the original criterion, while the comparison between mobile and one-rigid is unaffected, both arms being scored the same way.

The main metric is the *unique success rate* per task, *U/*200: Foldseek [25] clusters the generated design backbones among the PMPNN-final successful designs only (TM-score [36] cutoff 0.5), separately within each task. The raw yield *S/*200 is reported alongside it in the supplement (Figures S10 and S11). A task counts as solved if *U* ≥1.

Under the same protocol we ran the original ProteinZen on the benchmark tasks and on the original Protpardelle tasks [22] from which they were derived. These numbers set a reference point — the level of success of the base model in its own setting; the shift produced by finetuning and by motif mobility during generation is read against it. The run also confirms that the criterion and the Foldseek clustering behave as expected on a familiar set. The values in Figures 4 and 5 are computed over the tasks of our benchmark.

## Author Contributions

**I.P**.: Conceptualization, Methodology, Software, Investigation, Formal analysis, Data curation, Validation, Visualization, Writing – original draft. **E.G**.: Validation, Writing – review and editing. **A.G**.: Conceptualization, Supervision, Resources, Funding acquisition, Writing – review and editing.

### Acknowledgments

Computational resources were provided by the Faculty of Bioengineering and Bioinformatics, Lomonosov Moscow State University.

## Artificial Intelligence Disclosure

Large language model assistants were used to translate the manuscript from Russian into English, to edit the English text, to check bibliographic records, and, through an AI-assisted code editor, to help write and refactor parts of the software. The research design, the experiments and the interpretation of the results are the authors’ own. All text and code were reviewed, tested and verified by the authors, who take full responsibility for the content.

## Data Availability

NomadFlow is freely available at https://github.com/ipermyakov/nomadflow and is released under the MIT License; it derives from ProteinZen [19]. The repository contains the inference code, a pinned environment specification and a worked example that builds a scaffold for a multi-group task. The trained model weights are deposited at https://doi.org/10.5281/zenodo.22937971 under CC-BY-4.0. The definitions of the 21 benchmark tasks reported here and the source data behind every figure and table are included in the repository. The release covers inference; training code is not part of it.

## A Construction of multi-group training examples

### A.1 Why the construction is arranged this way

Two naive orderings break an example: (1) crop first, motif afterwards — the cropped fragment may have no room for the groups; (2) motif first on the full protein, then an independent crop — the crop may cut off part of a group that has already been chosen. Cells and candidate motifs are therefore built on the full protein, and a motif enters an example only together with **all** of its cell.

### A.2 Algorithm

1. **Partition into cells**. Per chain: the number of centers is *n* = ⌈*N*_chain_*/r*_target_⌉, chosen by farthest-point sampling over C*α*; residues go to the nearest center, subject to `cluster_max_res`.
2. **Candidate motif** (a candidate, not yet in the example) — a subset of each non-empty cell: either a compact multi-fragment active site (the catalytic type in the code), or the spatial core of the cell after erosion by Voronoi depth (the structural type). The proportion of such centers and their sizes are hyperparameters, listed below.
3. **Crop by whole cells** within the token budget (scaffold + copies of the motif + aggregators). A cell is accepted in full, and a motif is activated only together with it (with a probability, or under a quota on the number of groups). Sequence contact with the scaffold collected so far is required after the first cell.
4. **Topping up the scaffold** with individual residues, creating no motifs; then tokenization and one virtual aggregator per activated group.

**Invariant:** every activated motif is a subset of a cell accepted in full; the topping-up residues carry no group id; the number of tokens is ≤ `max_crop_residues`.

### A.3 Mixture and hyperparameters

The task mixture (FrameDiff [28] + AFDB [37]) consists of unconditional generation, indexed / unindexed mixed motif, and Voronoi unindexed multi-group. Both finetuning stages use the same mixture and differ only in crop size (`max_crop_residues` 224, then 320). The datasets are drawn with probabilities 0.25 (FrameDiff) and 0.75 (AFDB); within each, tasks are sampled in the same proportions: unconditional generation 0.10, indexed motif scaffolding 0.10, unindexed motif scaffolding 0.30, Voronoi unindexed multi-group 0.50. Half of all finetuning steps therefore fall on the multi-group regime, the regime for which the group is introduced.

The time *t* for the first three tasks is sampled from a lognormal (*µ* = 0, *σ* = 1), and for Voronoi multi-group **uniformly** on [*t*_min_, *t*_max_] = [0.01, 0.99]. Uniformity matters here: it is what justifies the factor 1*/t*_lock_ on the group terms (see §5.1.4) — without it the fraction of steps on which a group is still mobile equals *t*_lock_, and the contribution of those terms would be drowned out by the scaffold. The upper bound *t*_max_ = 0.99 *> t*_lock_ means that the post-lock phase is trained as well.

This algorithm builds training examples only. The benchmark partitions (`per_segment` / structural segments, active sites) are set separately, by contigs and `motif_groups` in the sampling configs.

## B Training hyperparameters

### B.1 Objective weights

The flow term enters with coefficient 1 and sets the scale against which the remaining weights are read.

Two refinements to the expression above. The rotation term is decomposed into an angular and an axial part, L_rot_ = 0.5 L_angle_ + L_axis_, rather than taken as a single norm. In L_seq_ seven residue types (CYS, GLU, HIS, PRO, GLN, ARG, TRP) enter with double weight. Both are inherited from ProteinZen and do not affect how the group terms are to be read.

### B.2 Optimization and the schedule of the stages

The starting point is the ProteinZen checkpoint `epoch-3278-step-780000`, loaded with `weights_only`, so the optimizer state is not carried over. In total, finetuning takes about 37 000 optimizer steps and on the order of 112 hours on four GPUs — against 780 000 steps of pretraining, that is, under five percent of it. The crop sizes by stage are 224 and then 320 residues (Table 2).

**Table 2.**
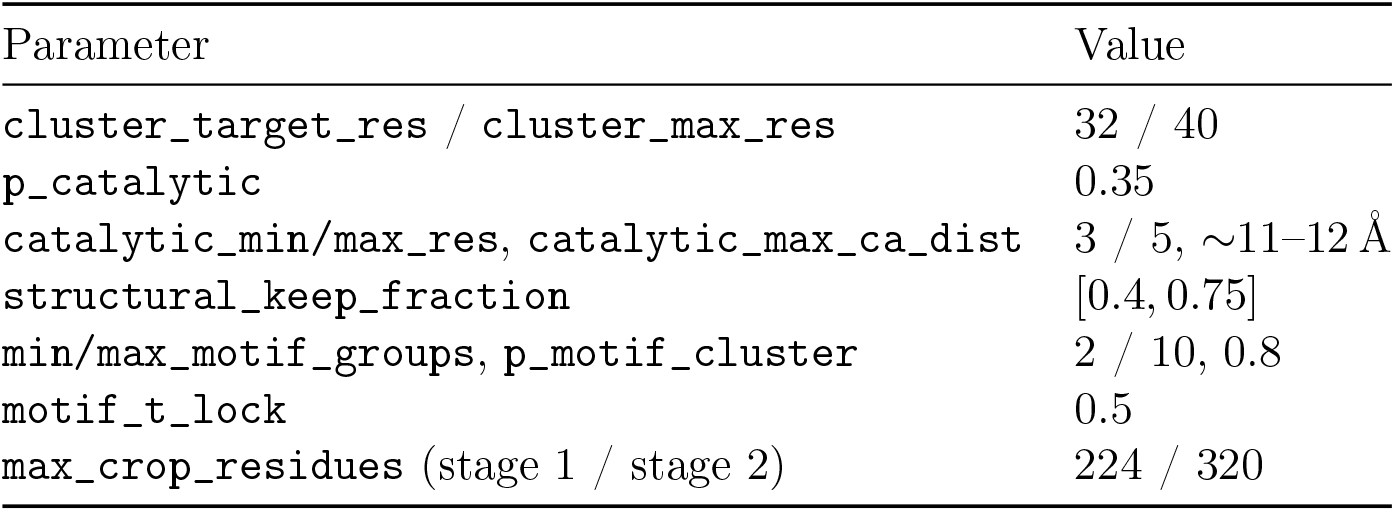
Hyperparameters of the construction of multi-group training examples.

| Parameter | Value |
| --- | --- |
| cluster_target_res / cluster_max_res | 32 / 40 |
| p_catalytic | 0.35 |
| catalytic_min/max_res, catalytic_max_ca_dist | 3 / 5, $\sim 11\text{--}12\text{ \AA}$ |
| structural_keep_fraction | [0.4, 0.75] |
| min/max_motif_groups, p_motif_cluster | 2 / 10, 0.8 |
| motif_t_lock | 0.5 |
| max_crop_residues (stage 1 / stage 2) | 224 / 320 |

**Table 3.** Weights of the terms of the objective. The values are the defaults of ProteinzenLModule(proteinzen/runtime/lmod.py); they are not overridden in the spatial / Voronoi finetuning.

| Term | parameter | weight |
| --- | --- | --- |
| Frame VF (trans + rot) | — | 1.0 |
| Aggregator pose, $\lambda_{\text{agg}}$ | `motif_aggregator_loss_weight` | 0.2 |
| Aggregator FAFE, $\lambda_{\text{FAFE}}^{\text{agg}}$ | `motif_aggregator_fafe_loss_weight` | 0.35 |
| Intergroup FAFE, $\lambda_{\text{FAFE}}^{\text{inter}}$ | `motif_intergroup_fafe_loss_weight` | 0.2 |
| Base FAFE, $\lambda_{\text{FAFE}}$ | `fafe_loss_weight` | 0.5 |
| Sequence, $\lambda_{\text{seq}}$ | `seq_loss_weight` | 0.25 |
| *internal reweightings, not terms of their own* |  |  |
| Aggregator frames in VF and FAFE, $w_{\text{agg}}$ | `motif_aggregator_weight` | $\times 5$ |
| Atomized rigids | `atom_rigid_upweight` | $\times 10$ |
| Angular part of $\mathcal{L}_{\text{rot}}$ | `rot_angle_weight` | 0.5 |
| Rare residue types in $\mathcal{L}_{\text{seq}}$ | `seq_weight` | $\times 2$ |

The crop224 stage runs for about 18 000 steps with gradient clipping 5.0, starting at a learning rate of 5 · 10^−4^ and following the step decay given below. The crop320 stage continues from its output for about 19 000 further steps at 5 · 10^−5^ with the same clipping.

The effective batch at every stage is 24 chains: four GPUs with gradient accumulation, which on crop224 is 2 ×3 per device and on crop320 1 × 6. The optimizer is Adam, *β* = (0.9, 0.999), *ε* = 10^−8^, weight decay 0. The learning rate is lowered in steps: from step 7000, every 3000 steps, by a factor of 0.5; cosine and warmup are switched off. Wall-clock time on a single node with four GPUs is about 40 hours for crop224 and about 72 for crop320.

## C Sampling with *t*_lock_

Methods fixes the local time of the groups and the fact that the group step is switched off after the lock. The underlying stochastic scaffold integrator (churn, *g*(*t*), score) is as in ProteinZen [19]; what follows concerns only how the aggregator step is reconciled with the two times.

### C.1 Integrator constants

In all runs reported here: a uniform grid of 400 steps from *t*_min_ = 10^−2^ to 1, with *t*_lock_ = 0.5. Churn with coefficient 0.4 — at every step the state is rolled back to 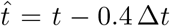 and noised afresh. Update step sizes of 1.5 for translations and rotations at *γ* = 0.16; the rotation schedule is linear, as is the schedule of *t*_motif_ . Centering of the system on the motif is switched on, so translations are everywhere measured from the motif part rather than from the center of mass of the whole chain. The prior on the group pose is *σ*_grp_ = 12 Å (Methods), and that of the scaffold 16 Å.

### C.2 Equations for the group step

A global grid *t* ∈ [0, 1] with step Δ*t*. The local time of the groups (linear schedule) is

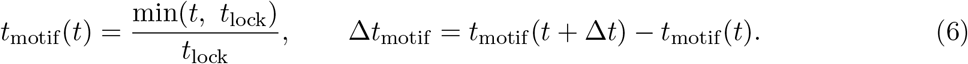

For *t ≥ t*_lock_: *t*_motif_ = 1 and the group step is off.

For the aggregator the velocity field is evaluated in the local time, as in training:

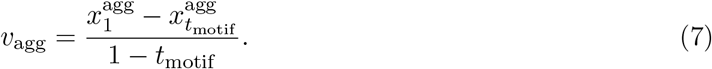

The Euler translation step on the global grid is rescaled back to Δ*t*:

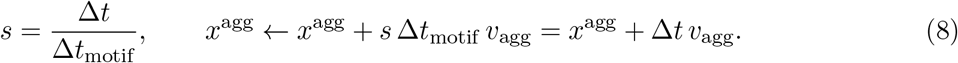

Before the lock, with a linear schedule, *s* = *t*_lock_. Without *s* the dynamics of the groups would part company with the parameterization used in training: the velocity field lives in *t*_motif_ while the grid step is Δ*t*.

The SE(3) delta taken from the updated aggregator is applied rigidly to all frames of the group (`motif_group_euler_step`). The scaffold continues to be integrated by churn and Euler in the global (*t*, Δ*t*).

#### Churn and the rigidity of a group

The ProteinZen per-rigid churn is switched off on the **real** motif frames: independent noise per residue would pull the prescribed geometry apart. The noise on a group is a single common rigid SE(3) churn (a delta on the aggregator), applied by the same transformation to the whole group (`motif_group_churn`). The scaffold is noised residue by residue, as in the original integrator.

### C.3 Pseudocode for a step (before the lock)

At a global step *t* → *t* + Δ*t*:

1. Scaffold: per-rigid churn / Euler in the global (*t*, Δ*t*) (ProteinZen).
2. Aggregator: group-level churn (a single SE(3) delta) →`motif_group_churn` onto all frames of the group; per-rigid churn on the real motif is off.
3. Aggregator Euler: (*t*_motif_, Δ*t*_motif_), *v*_agg_, and the shift with scale *s* (equations above).
4. `motif_group_euler_step`: carry the whole group by the aggregator delta.
5. For *t* ≥ *t*_lock_: steps 2–4 are skipped and the motif is fixed conditioning.

## Supporting Information

This section holds additional **results**: the composition of the tasks and the native structures of those same tasks (Figures S1–S2, immediately after Tables S1–S2), a per-task table of *S* and *U* under the main protocol (ProteinMPNN + ESMFold), and quantitative panels accompanying the figures of the main text.

### Native structural segments

**Figure S1.**
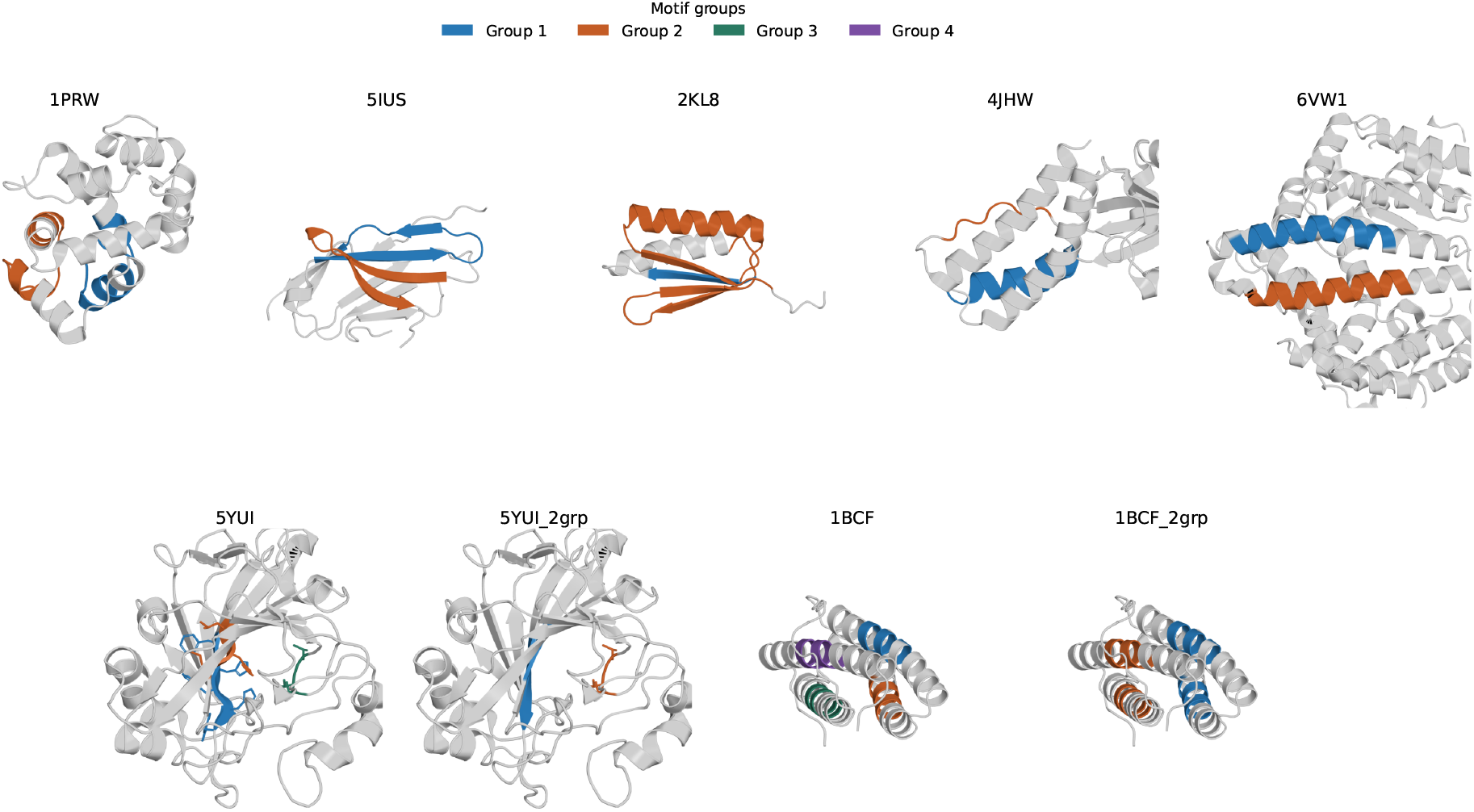
Native structures of the tasks on the structural segments axis. The scaffold is gray; the colors are the motif groups from the sampling configs, in the order of Table S1. The seven proteins are shown in the full split, each group one fragment; 5YUI and 1BCF are accompanied by `_2grp`, the same set of residues fused into two groups. The camera is shared within a PDB /`_2grp` pair. These are task definitions, not design results.

### Native active sites with additions

**Figure S2.**
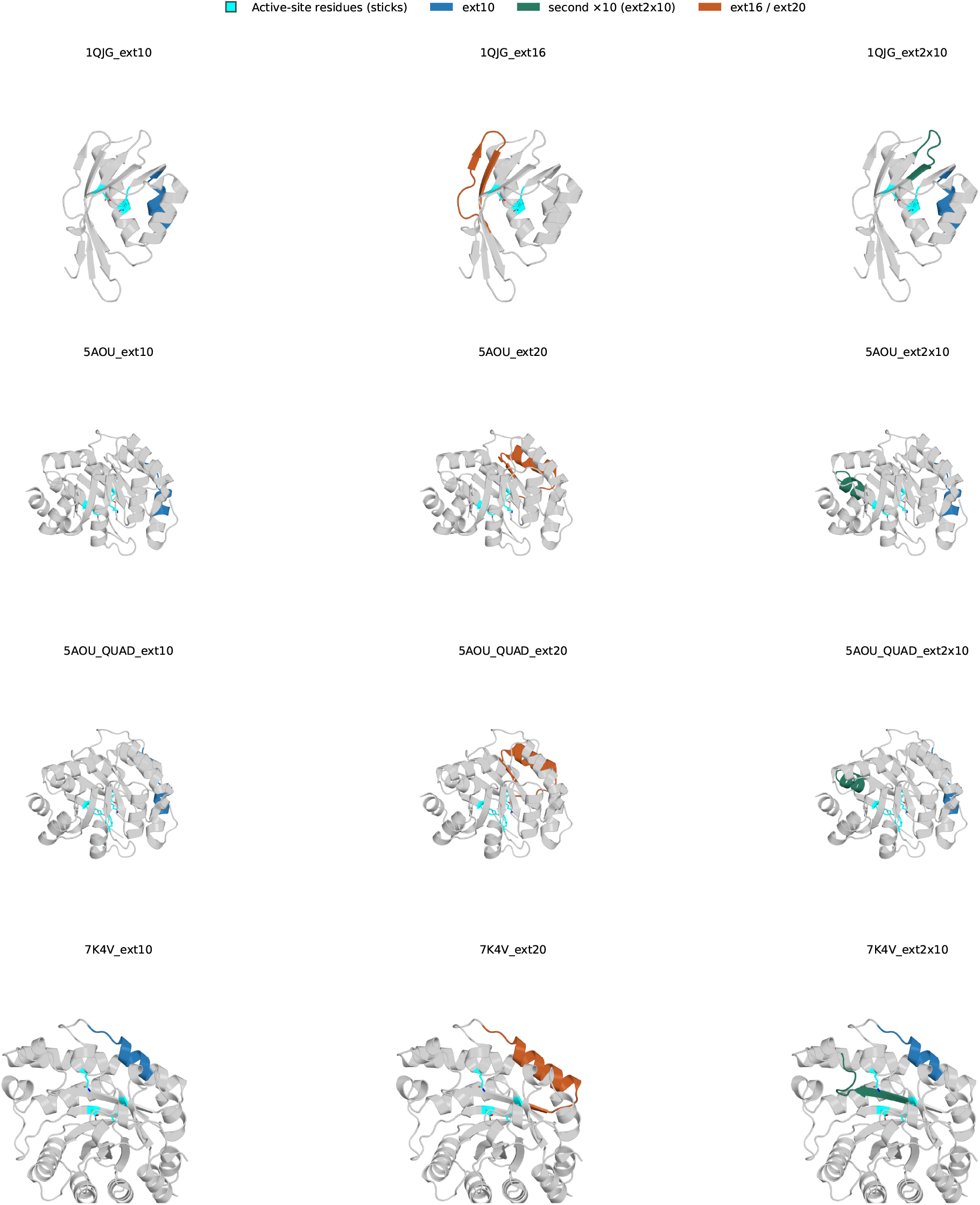
Native structures of the active-site tasks with additions. Each cell is a separate task, not a superposition of the additions: the active site as sticks (cyan) plus the segments of that task alone. Rows are the families 1QJG, 5AOU, 5AOU_QUAD and 7K4V, with the camera shared along a row; columns are `ext10`, the lengthening (`ext16` for 1QJG, `ext20` for the rest) and ext2x10. These are task definitions, not design results.

### Composition of the benchmark tasks

The partitions come from the configs the results were computed with: `configs_multi/config_ multi_motif_se3_per_segment.yaml` and `configs_multi/config_multi_motif_se3_catalytic_ tlock_0.5.yaml`; the `ext0` tasks come from `configs_multi/config.yaml`. *F* is the number of fragments (a continuous contig, or a single residue of an active site), *m* the number of independent rigid groups in the multi-group partition, *m ™* 1 the number of free mutual poses, *n* the number of motif residues, and *L* the length of the generated chain (`total_length` in the config). Parentheses give the size of a group. Braces mark several fragments bound into one group. One-rigid uses the same set of residues as a single group (*m* = 1) and the same *L*.

The length of each design is sampled uniformly from the range *L*, independently for each of the 200 designs of a task (for 2KL8 the length is fixed). The ranges are inherited from the original Protpardelle tasks and shifted by the length of the added segments, so both moves on the active-site axis — lengthening and adding a fragment — spend the same scaffold budget: within an `ext20 / ext2x10` pair the range of *L* coincides, and all that differs is what the added residues are invested in. A machine-readable copy is at `article_results/figures/publication/source_data/si_task_ composition.csv`. The native structures of the same partitions are shown in Figure S1 (structural segments) and Figure S2 (active sites with additions).

**Table S1.** Composition of the benchmark tasks: the structural segments axis (Figure 4).

| Task | $F$ | $m$ | $m - 1$ | $n$ | $L$ | group composition |
| --- | --- | --- | --- | --- | --- | --- |
| 1PRW | 2 | 2 | 1 | 40 | 60–150 | A16-35 (20); A52-71 (20) |
| 5IUS | 2 | 2 | 1 | 42 | 57–142 | A119-140 (22); A63-82 (20) |
| 2KL8 | 2 | 2 | 1 | 59 | 79 | A1-7 (7); A28-79 (52) |
| 4JHW | 2 | 2 | 1 | 24 | 60–90 | F196-212 (17); F63-69 (7) |
| 6VW1 | 2 | 2 | 1 | 38 | 62–83 | A24-42 (19); A64-82 (19) |
| 5YUI | 3 | 3 | 2 | 11 | 50–100 | A93-97 (5); A118-120 (3); A198-200 (3) |
| 5YUI_2grp | 3 | 2 | 1 | 11 | 50–100 | {A93-97, A118-120} (8); A198-200 (3) |
| 1BCF | 4 | 4 | 3 | 32 | 96–152 | A92-99 (8); A123-130 (8); A47-54 (8); A18-25 (8) |
| 1BCF_2grp | 4 | 2 | 1 | 32 | 96–152 | {A92-99, A123-130} (16); {A47-54, A18-25} (16) |

**Table S2.**
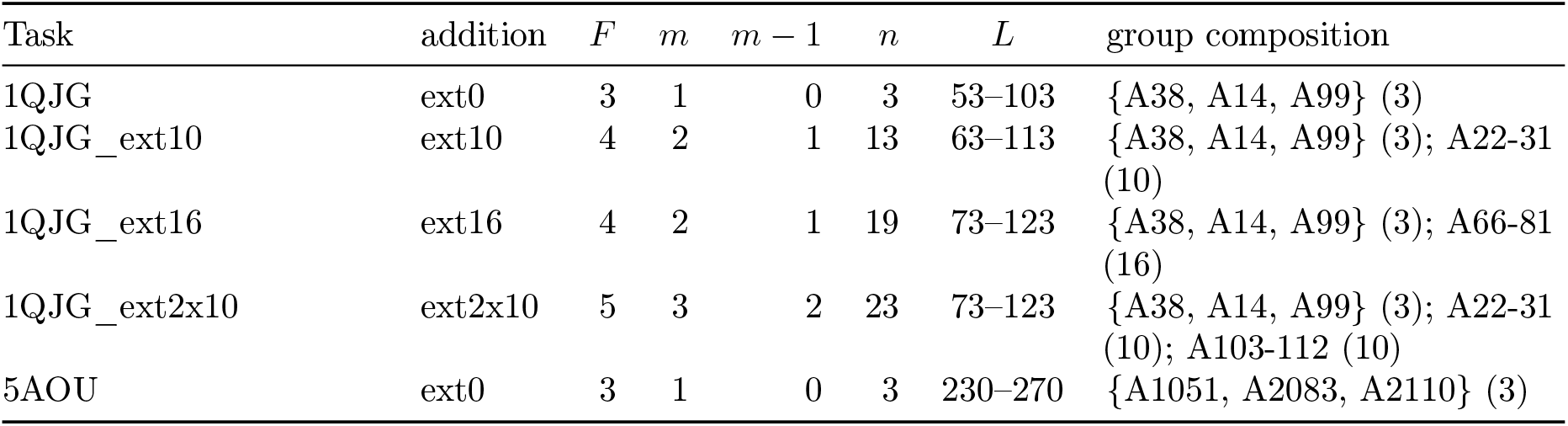
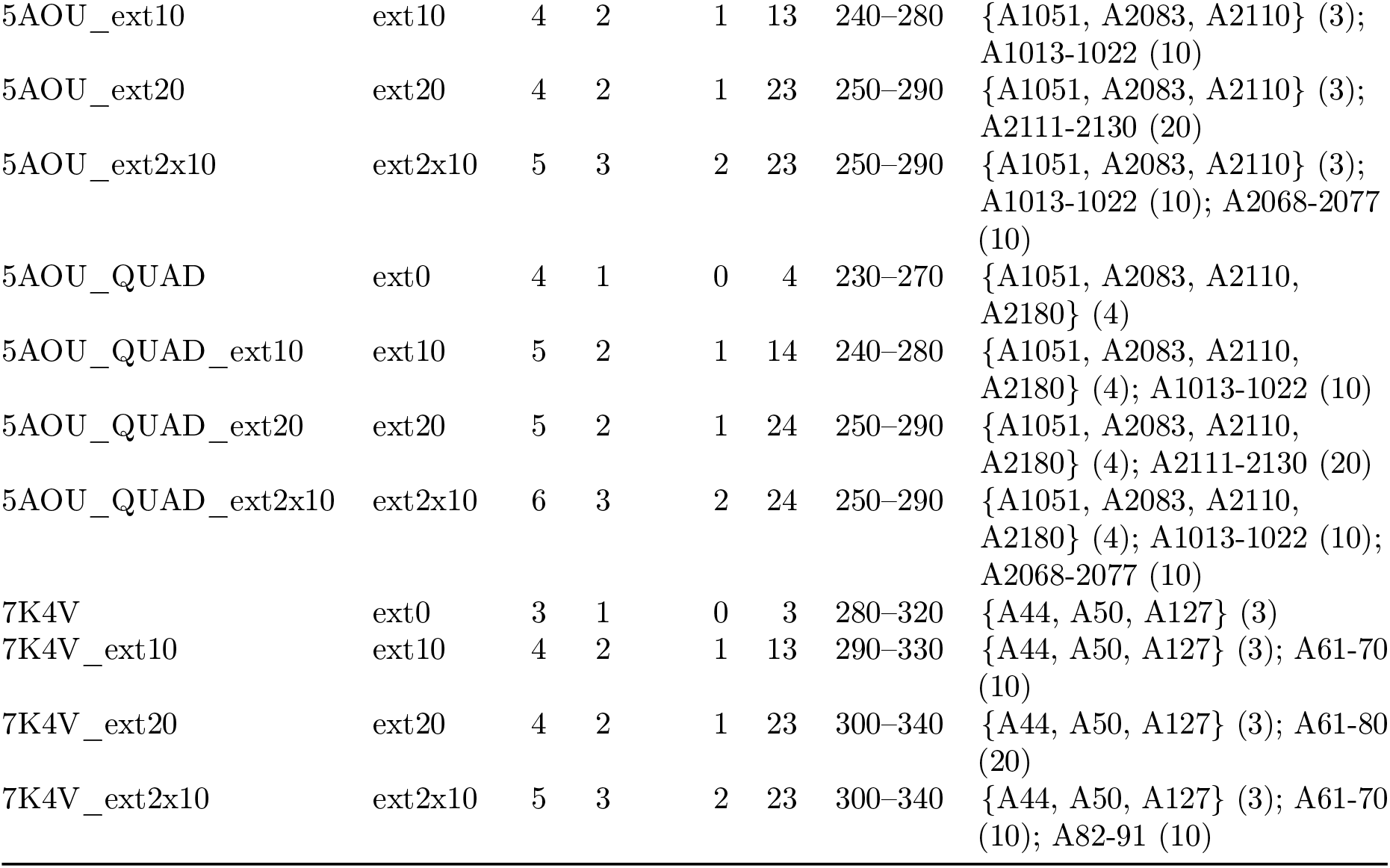
Composition of the benchmark tasks: the active-site axis (Figure 5). Tasks with no ext* suffix are ext0, one-rigid only.

### Per-task *S* **and** *U*

Every row has *N* = 200. The raw yield *S* is the number of designs with at least one PMPNN sequence that reproduced all groups, and unique *U* the number of Foldseek clusters (TM 0.5) among those designs, for the same series as in Figures 4–5, S10 and S11: one-rigid, mobile and `_2grp`. A machine-readable copy is at `article_results/figures/publication/source_data/si_pmpnn_ task_table.csv`. One-shot does not enter this table (Figures S8–S9).

**Table S3.** Per-task *S* and *U* (ProteinMPNN + ESMFold): the structural segments axis.

| Task | Series | $N$ | $S$ | $U$ |
| --- | --- | --- | --- | --- |
| 1PRW | one-rigid | 200 | 117 | 33 |
| 1PRW | mobile | 200 | 153 | 92 |
| 5IUS | one-rigid | 200 | 3 | 2 |
| 5IUS | mobile | 200 | 2 | 1 |
| 2KL8 | one-rigid | 200 | 41 | 2 |
| 2KL8 | mobile | 200 | 18 | 2 |
| 4JHW | one-rigid | 200 | 3 | 3 |
| 4JHW | mobile | 200 | 0 | 0 |
| 6VW1 | one-rigid | 200 | 182 | 34 |
| 6VW1 | mobile | 200 | 160 | 43 |
| 5YUI | one-rigid | 200 | 94 | 87 |
| 5YUI | mobile | 200 | 114 | 112 |
| 5YUI_2grp | 2grp | 200 | 101 | 93 |
| 1BCF | one-rigid | 200 | 184 | 27 |
| 1BCF | mobile | 200 | 185 | 49 |
| 1BCF_2grp | 2grp | 200 | 121 | 32 |

**Table S4.** Per-task *S* and *U* (ProteinMPNN + ESMFold): the active-site axis. The ext0 tasks are one-rigid only; for 1QJG the long addition is ext16.

| Task | Series | $N$ | $S$ | $U$ |
| --- | --- | --- | --- | --- |
| 1QJG (**ext0**) | one-rigid | 200 | 121 | 89 |
| 1QJG_ext10 | one-rigid | 200 | 64 | 36 |
| 1QJG_ext10 | mobile | 200 | 40 | 39 |
| 1QJG_ext16 | one-rigid | 200 | 63 | 22 |
| 1QJG_ext16 | mobile | 200 | 12 | 10 |
| 1QJG_ext2x10 | one-rigid | 200 | 62 | 26 |
| 1QJG_ext2x10 | mobile | 200 | 19 | 19 |
| 5AOU (**ext0**) | one-rigid | 200 | 16 | 7 |
| 5AOU_ext10 | one-rigid | 200 | 14 | 9 |
| 5AOU_ext10 | mobile | 200 | 42 | 22 |
| 5AOU_ext20 | one-rigid | 200 | 187 | 13 |
| 5AOU_ext20 | mobile | 200 | 143 | 13 |
| 5AOU_ext2x10 | one-rigid | 200 | 68 | 18 |
| 5AOU_ext2x10 | mobile | 200 | 3 | 2 |
| 5AOU_QUAD (**ext0**) | one-rigid | 200 | 5 | 3 |
| 5AOU_QUAD_ext10 | one-rigid | 200 | 2 | 2 |
| 5AOU_QUAD_ext10 | mobile | 200 | 12 | 4 |
| 5AOU_QUAD_ext20 | one-rigid | 200 | 150 | 7 |
| 5AOU_QUAD_ext20 | mobile | 200 | 77 | 7 |
| 5AOU_QUAD_ext2x10 | one-rigid | 200 | 42 | 7 |
| 5AOU_QUAD_ext2x10 | mobile | 200 | 3 | 2 |
| 7K4V (**ext0**) | one-rigid | 200 | 34 | 16 |
| 7K4V_ext10 | one-rigid | 200 | 12 | 4 |
| 7K4V_ext10 | mobile | 200 | 2 | 2 |
| 7K4V_ext20 | one-rigid | 200 | 49 | 8 |
| 7K4V_ext20 | mobile | 200 | 15 | 5 |
| 7K4V_ext2x10 | one-rigid | 200 | 0 | 0 |
| 7K4V_ext2x10 | mobile | 200 | 0 | 0 |

### Pairwise density across the structural tasks

**Figure S3.**
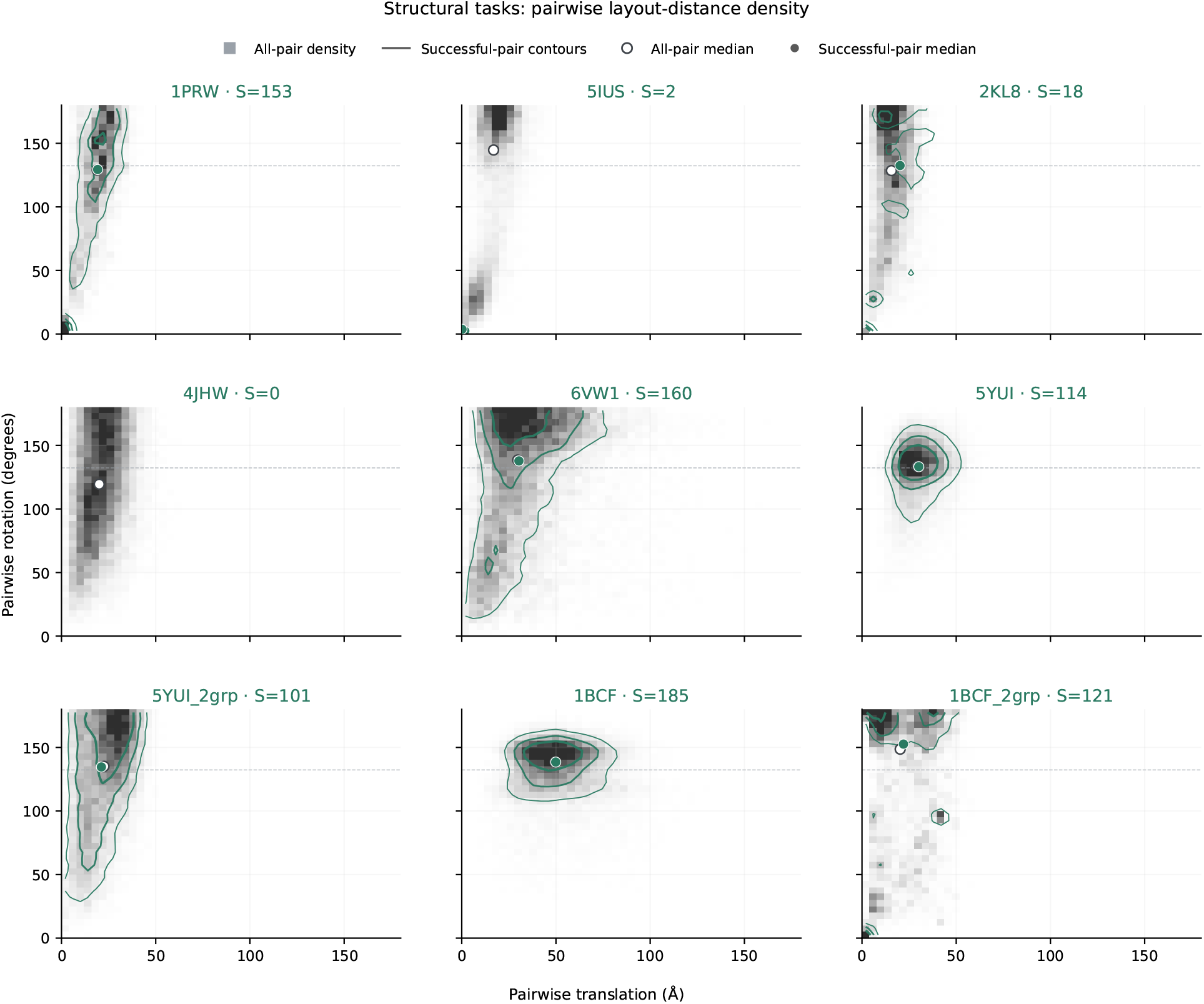
Joint distribution of pairwise distances between NomadFlow layouts across the nine structural tasks: translation (Å) against rotation (degrees). The gray background is the normalized density over all pairs of designs; the colored contours are pairs of designs that passed the PMPNN+ESMFold criterion. The open and filled markers are the medians of each coordinate for the two sets. Where successful pairs are few, the occupied density cells are shown instead of a contour. *S* in each panel title is the number of successful designs; the dashed line is 132.3^°^, the median angle between random rotations in SO(3) (Figure S4). Pairs within a task are dependent and are used to describe the shape of the distribution, not as independent observations.

### Summary of excursion and pairwise medians

**Figure S4.**
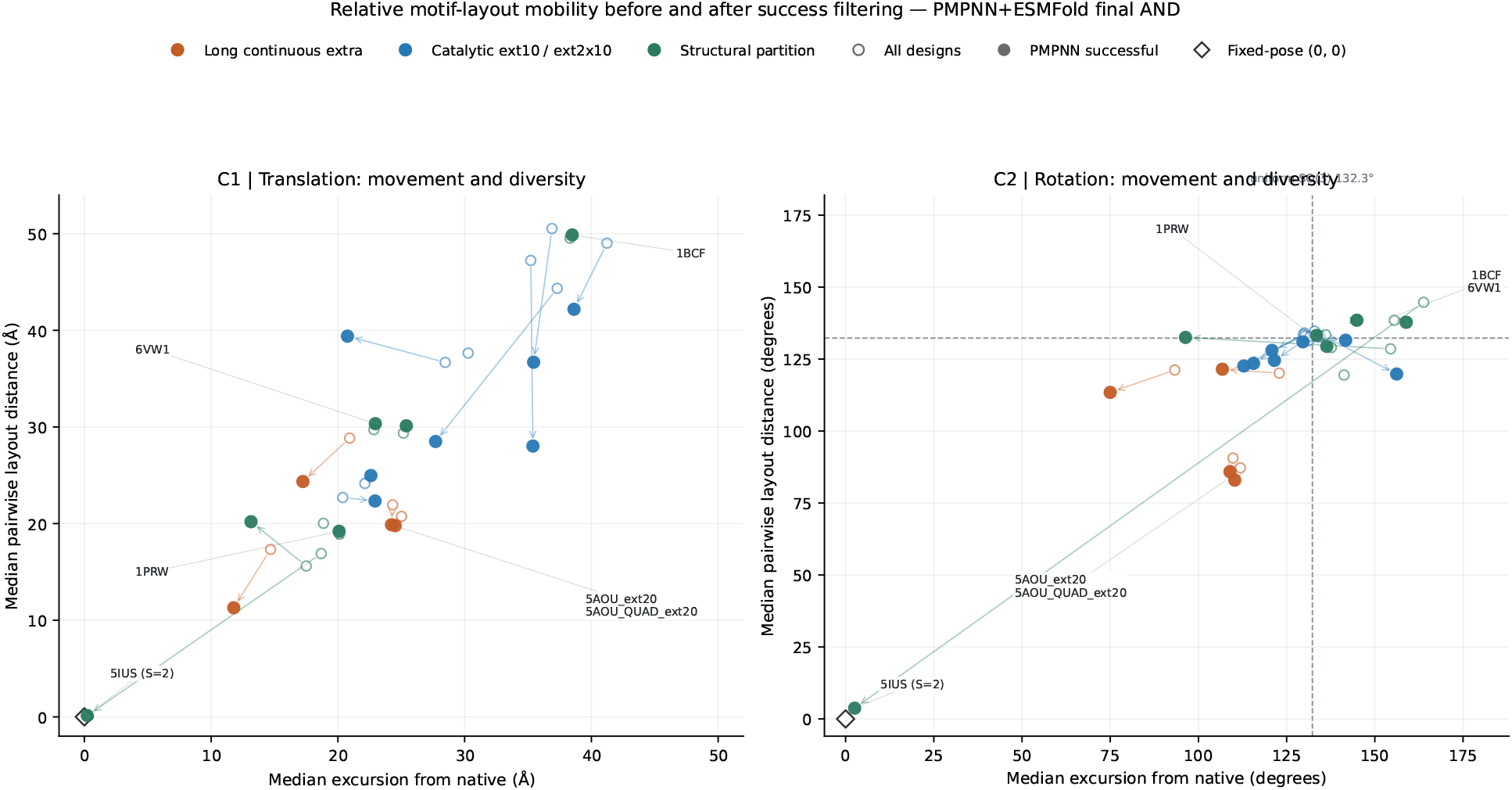
Median excursion from native against the median pairwise distance between designs, separately for translation (Å) and rotation (degrees). An open marker is all 200 NomadFlow designs; a filled one, those that passed the PMPNN+ESMFold criterion; the arrow is the step between them. Color indicates the type of condition. The diamond is the theoretical fixed-pose point (0, 0), not a measured design; ProteinZen / one-rigid is not shown. The dashed line in panel C2 (rotation) is 132.3^°^. the median angle between two independent, uniformly random rotations in SO(3), that is, the level reached by mutual orientations with no agreement between them whatsoever. Values at that line mean that in rotation the set of layouts is indistinguishable from random; values well below it mean that some agreement has been retained. This summarizes the medians of Figures 6–7; it shows neither the shape of the set nor whether native is visited by some of the designs.

### Pairwise density across the active-site tasks

**Figure S5.**
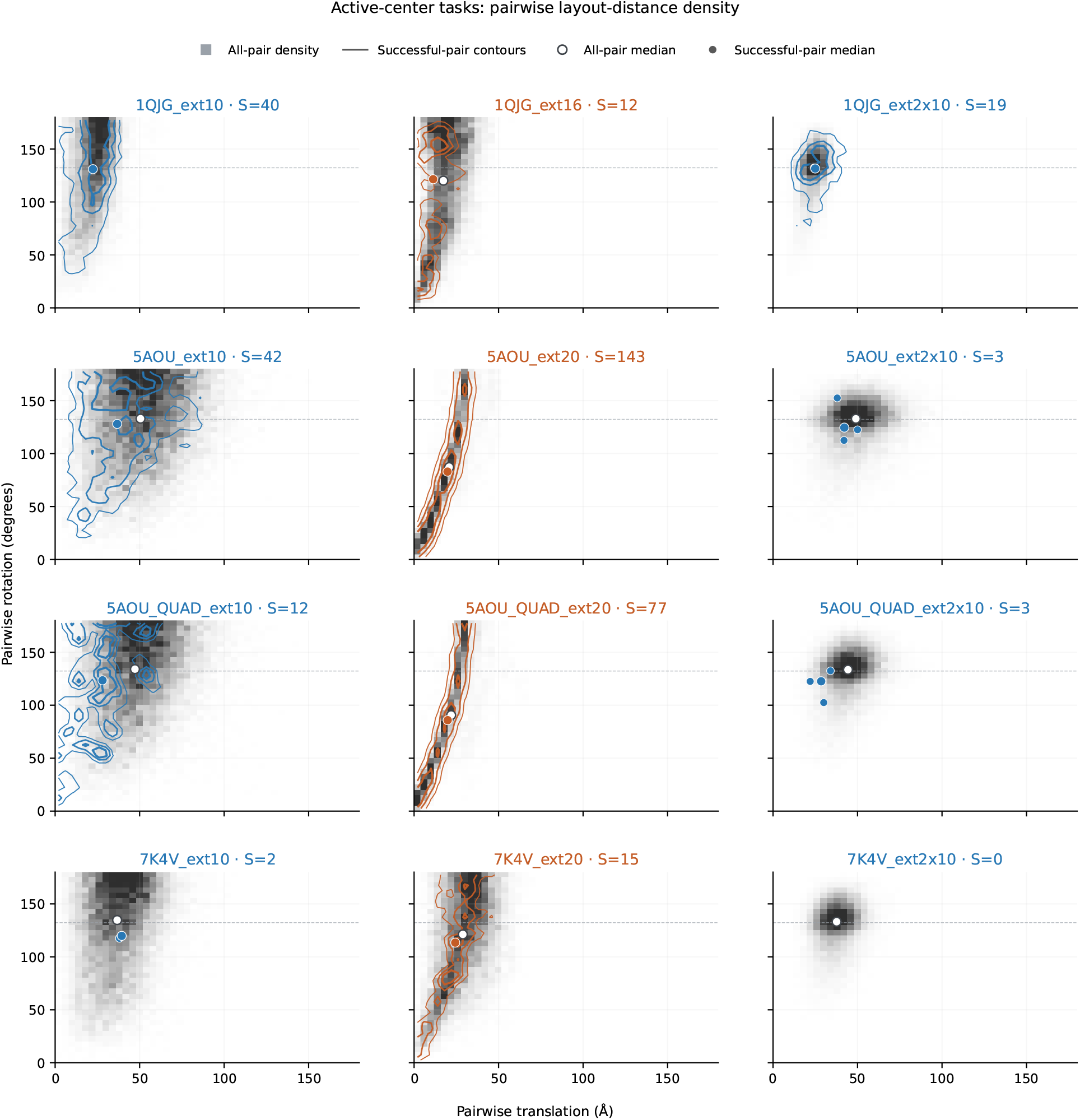
Joint distribution of pairwise distances across the twelve tasks with active sites and structural additions. Symbols as in Figure S3; the columns correspond to `ext10`, the long continuous segment and `ext2x10`.

### Native-anchored MDS across the structural tasks

**Figure S6.**
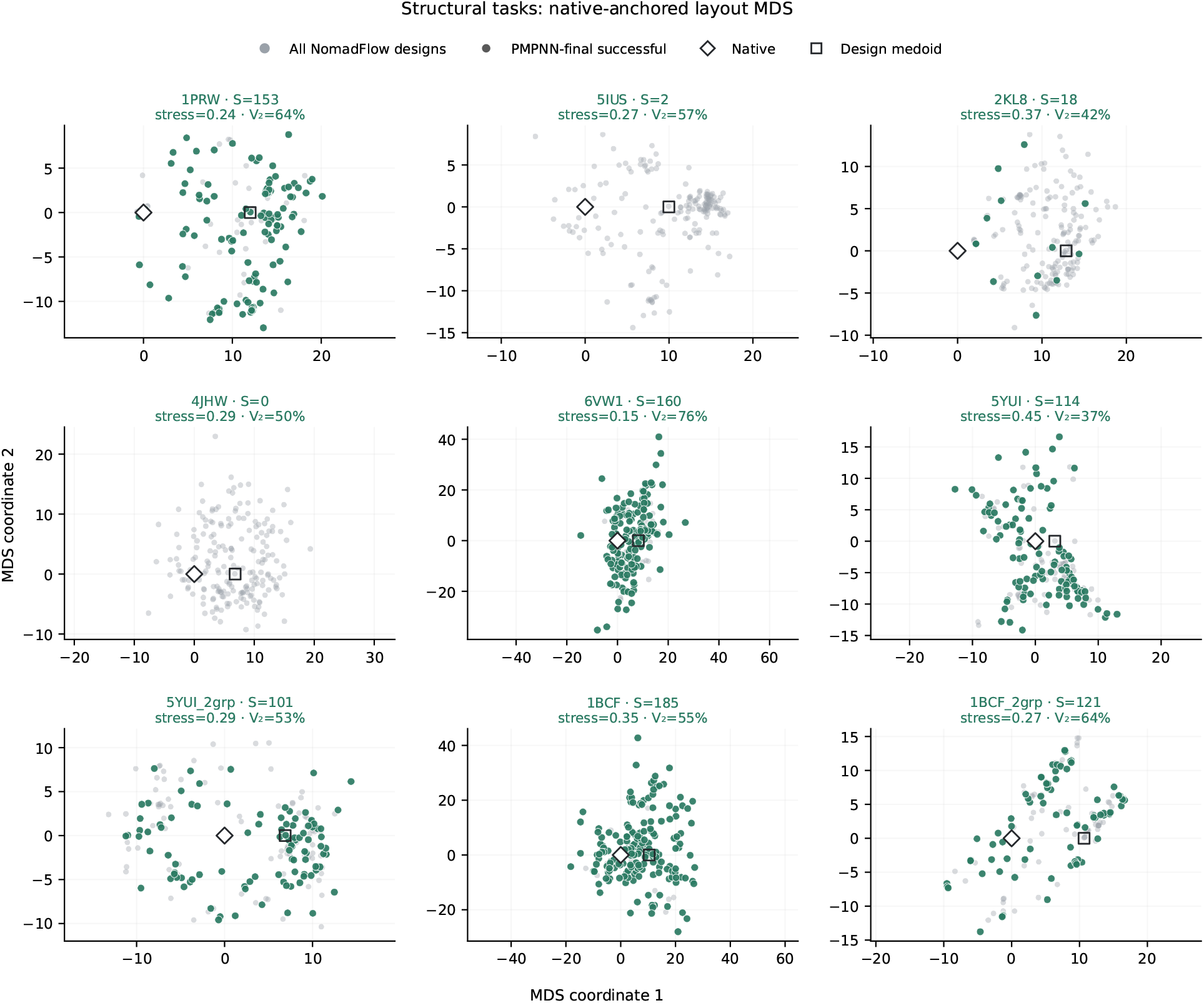
Two-dimensional classical MDS of all 200 NomadFlow layouts together with native, for each structural task. The distance combines the pairwise differences of the relative translations and rotations at scales of 2 Å and 15^°^. Native is moved to the origin, and the orientation of the axes is chosen so that the medoid of the designs lies on the positive first axis. Gray points are all designs, colored ones those that passed the PMPNN+ESMFold criterion, and the square is the medoid. Panel titles give the Kruskal stress and the fraction of the positive spectrum retained by the two coordinates *V*_2_. The MDS axes carry no physical meaning of their own.

### Design→fold retention

**Figure S7.**
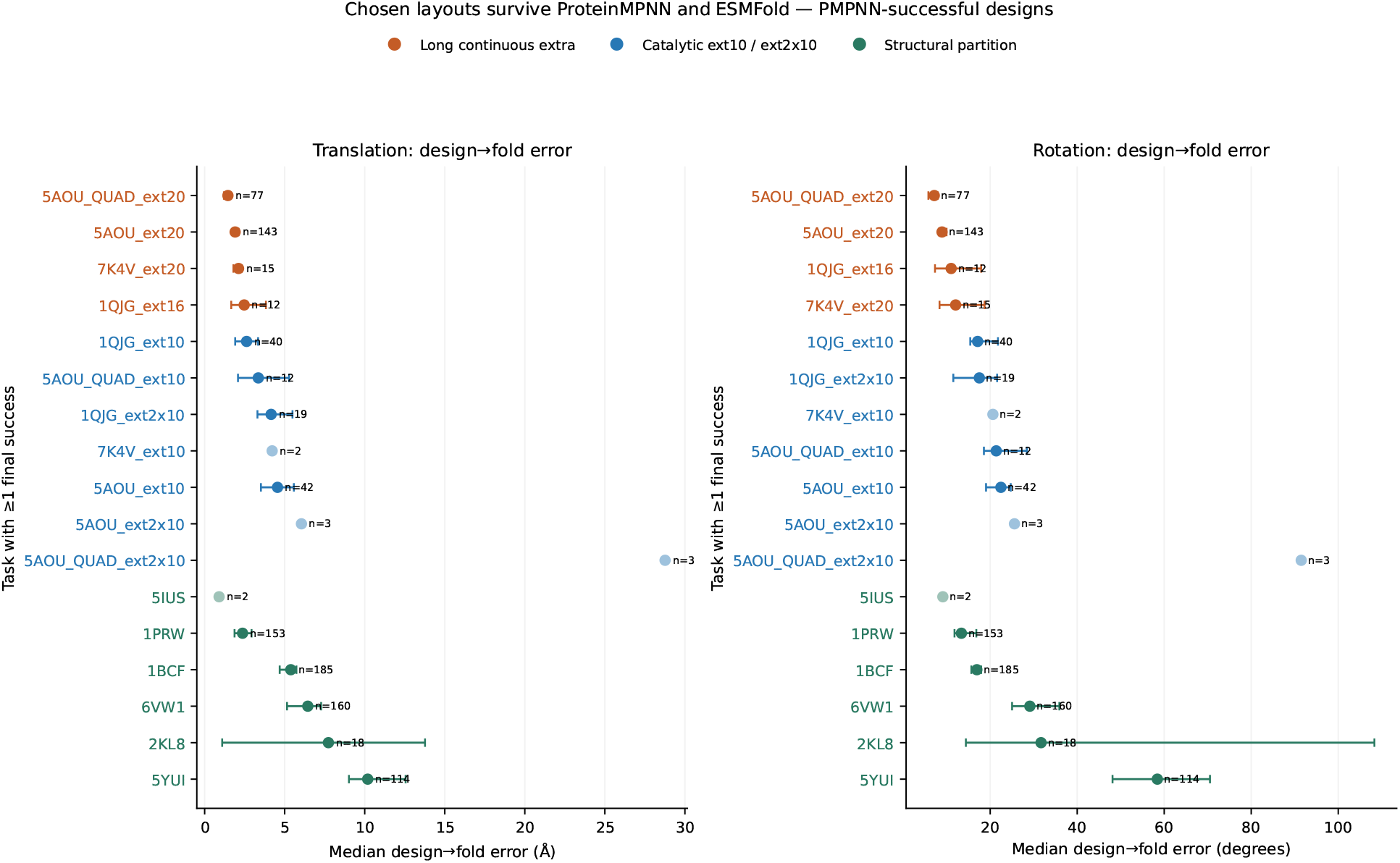
Median design fold → error among the successful NomadFlow designs after ProteinMPNN and ESMFold, separately for translation (Å) and rotation (degrees). Color indicates the same type of condition as in Figure S4. Labels give *n*, the number of successful designs; no intervals are shown for *n <* 5. Muted markers are tasks with *n* ≤ 6. A check that the layouts of Figures 6–7 are largely retained through folding.

### One-shot unique, structural segments

**Figure S8.**
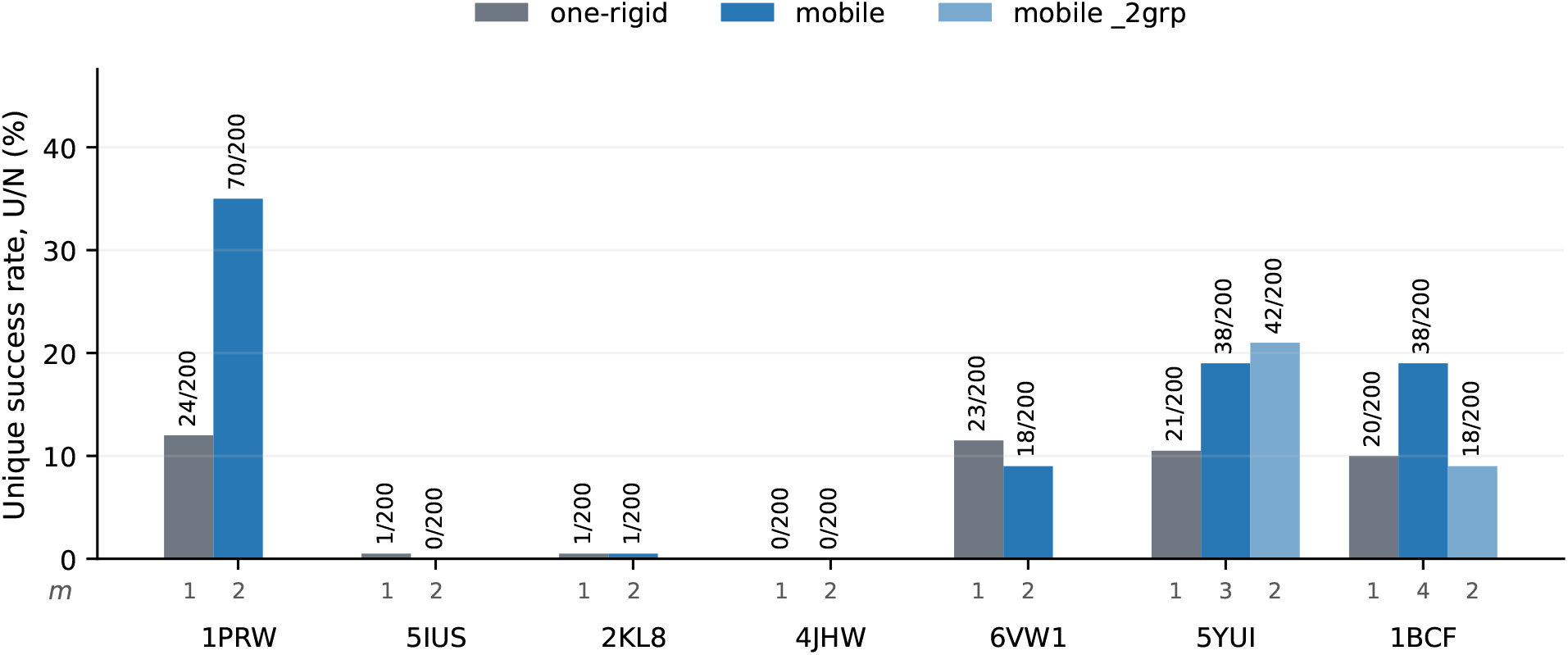
Unique success rate *U/*200 under the one-shot all-atom self-consistency protocol (without ProteinMPNN); the same proteins, residue sets, series and success criterion as in Figure 4. Foldseek clusters the successful design backbones among the one-shot successes. *N* = 200.

Without ProteinMPNN the same criterion makes success far rarer: on structural segments many tasks give *U* = 0. The panel is included as a self-consistency control on the protocol.

### One-shot unique, active sites

**Figure S9.**
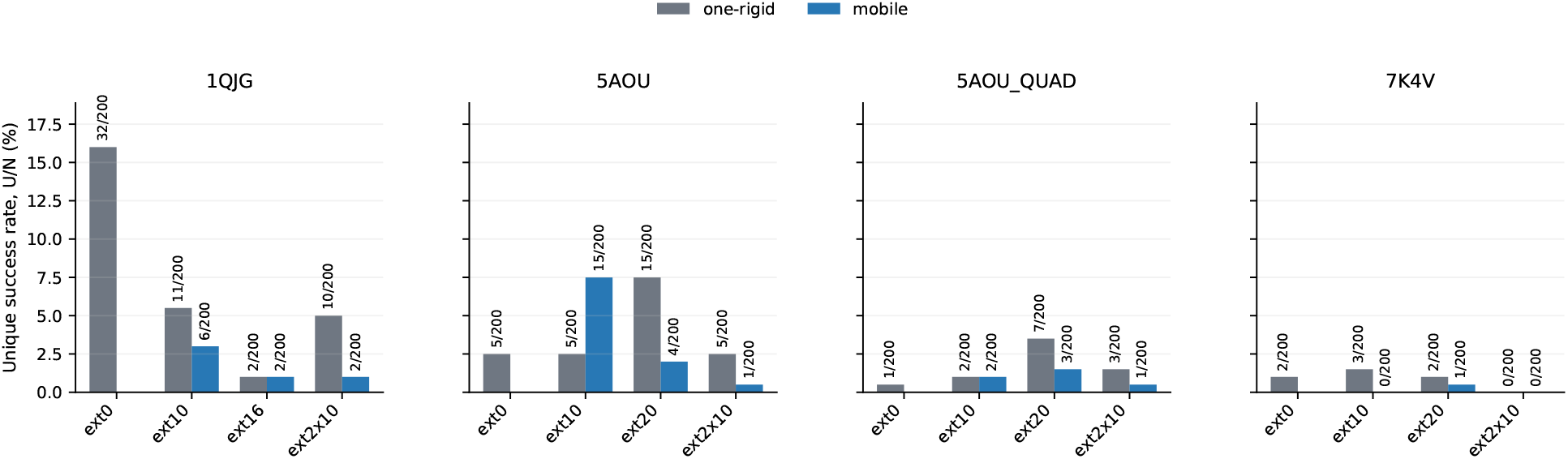
Unique success rate *U/*200 under one-shot self-consistency; the same families, additions and success criterion as in Figure 5. ext0 is one-rigid only. For 1QJG the long addition is `ext16`. *N* = 200.

7K4V does not go to zero entirely under one-shot (ext0 *U* = 2, ext20 NomadFlow *U* = 1). These numbers are not comparable with the PMPNN results of Figure 5.

### PMPNN raw yield, structural segments

**Figure S10.**
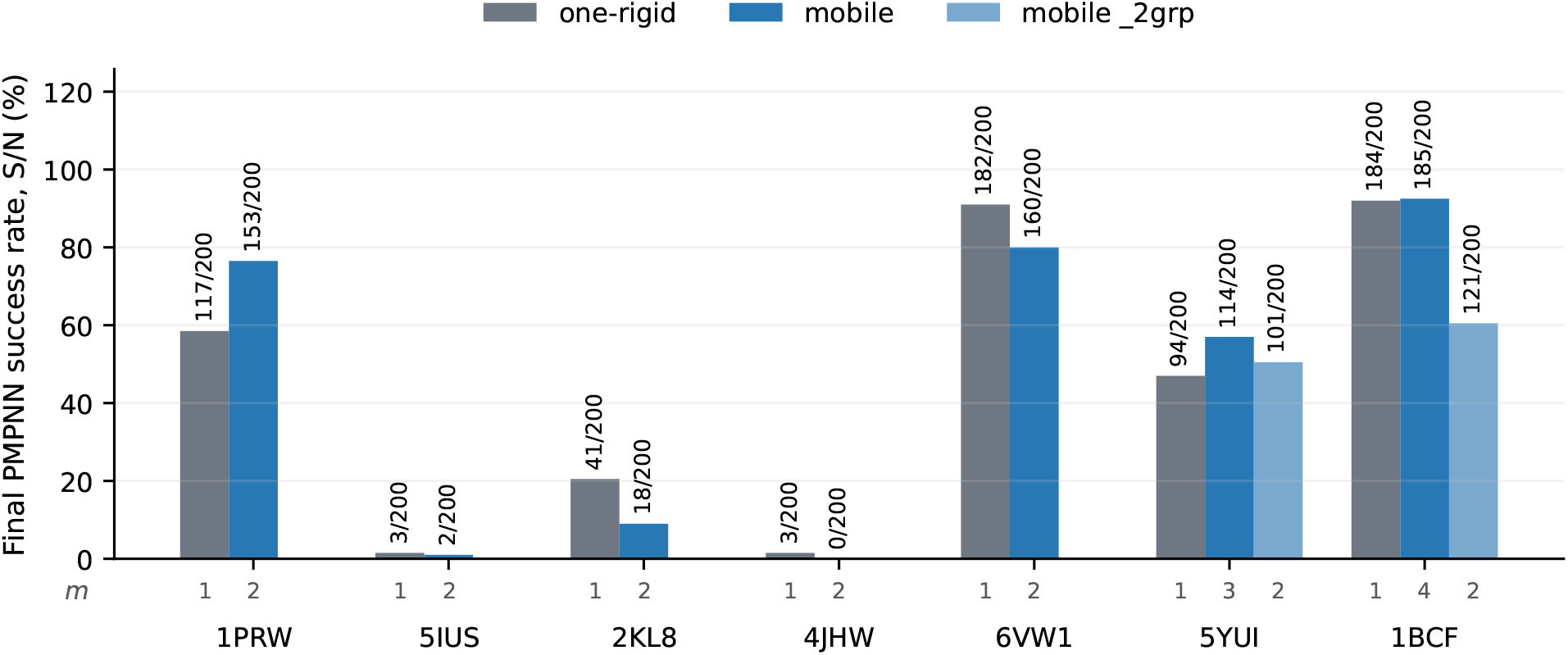
Raw PMPNN+ESMFold yield *S/*200 for the same seven proteins, residue sets, series and success criterion as in Figure 4. The figures beneath the bars are the number of groups *m*, as in Figure 4. Bar labels are *S/*200. Unique stays in Figure 4; this panel shows yield only. *N* = 200.

This shows how unique and yield come apart (6VW1: *S* 182 →160 while *U* 34 →43; 1BCF one-rigid nearly saturates yield at a moderate unique).

### PMPNN raw yield, active sites

**Figure S11.**
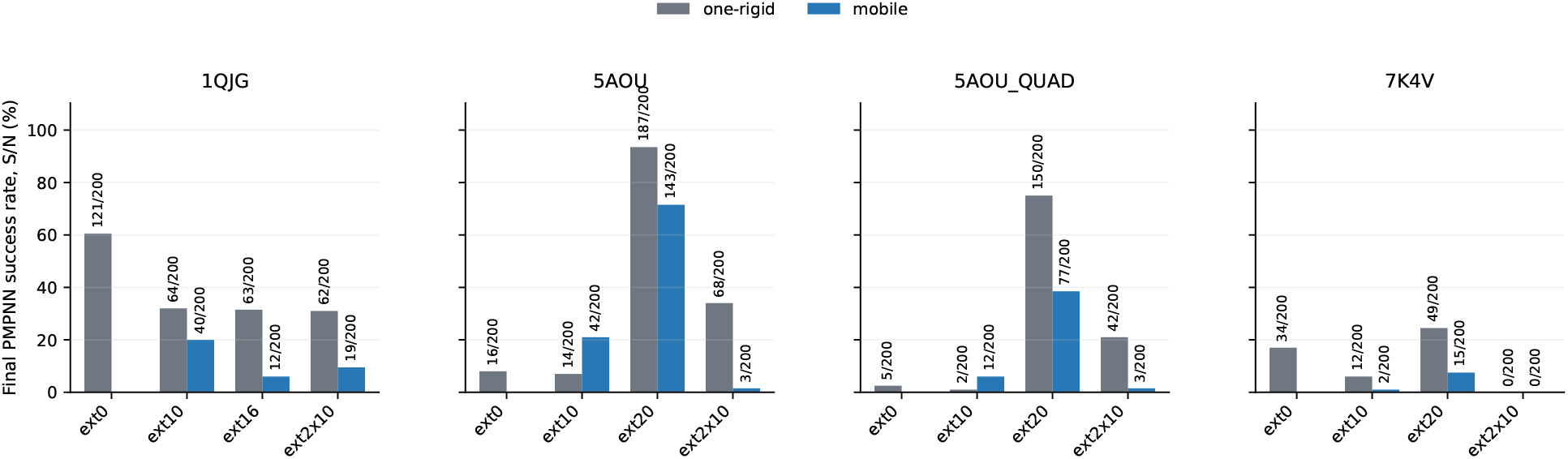
Raw PMPNN+ESMFold yield *S/*200 for the same active-site families, addition categories and success criterion as in Figure 5. ext0 is one-rigid only. For 1QJG the long addition is ext16. Bar labels are *S/*200. *N* = 200.

This is the picture behind the one-rigid collapse on 5AOU `ext20` (*S* = 187, *U* = 13) and QUAD `ext20` (*S* = 150, *U* = 7): a high yield does not mean an easy task.

## References

[1] Tanja Kortemme. De novo protein design—from new structures to programmable functions. Cell, 187(3):526–544, 2024. doi: 10.1016/j.cell.2023.12.028.

[2] Bruno E. Correia, John T. Bates, Rebecca J. Loomis, Gretchen Baneyx, Chris Carrico, Joseph G. Jardine, et al. Proof of principle for epitope-focused vaccine design. Nature, 507:201–206, 2014. doi: 10.1038/nature12966.

[3] Melissa L. Zastrow, Anna F. A. Peacock, Jeanne A. Stuckey, and Vincent L. Pecoraro. Hydrolytic catalysis and structural stabilization in a designed metalloprotein. Nature Chemistry, 4(2): 118–123, 2012. doi: 10.1038/nchem.1201.

[4] Markus Braun, Adrian Tripp, Morakot Chakatok, Sigrid Kaltenbrunner, Celina Fischer, David Stoll, Aleksandar Bijelic, Wael Elaily, Massimo G. Totaro, Melanie Moser, Shlomo Y. Hoch, Horst Lechner, Federico Rossi, Matteo Aleotti, Mélanie Hall, and Gustav Oberdorfer. Computational enzyme design by catalytic motif scaffolding. Nature, 649(8095):237–245, 2026. doi: 10.1038/s41586-025-09747-9.

[5] Anna Lauko, Samuel J. Pellock, Kiera H. Sumida, Ivan Anishchenko, David Juergens, Woody Ahern, et al. Computational design of serine hydrolases. Science, 388(6744):eadu2454, 2025. doi: 10.1126/science.adu2454.

[6] Yannan Qu, Chentong Wang, Hongli Zhu, Yanjun Wang, and Longxing Cao. De novo design of metalloproteases for targeted amyloid-β cleavage. bioRxiv preprint, 2026. Preprint posted January 6, 2026.

[7] Jue Wang, Sidney Lisanza, David Juergens, Doug Tischer, Joseph L. Watson, et al. Scaffolding protein functional sites using deep learning. Science, 377(6604):387–394, 2022. doi: 10.1126/science.abn2100.

[8] Joseph L. Watson, David Juergens, Nathaniel R. Bennett, Brian L. Trippe, Jason Yim, Helen E. Eisenach, Woody Ahern, et al. De novo design of protein structure and function with RFdiffusion. Nature, 620:1089–1100, 2023. doi: 10.1038/s41586-023-06415-8. URL https://www.nature.com/articles/s41586-023-06415-8.

[9] John B. Ingraham, Max Baranov, Zak Costello, Karl W. Barber, Wujie Wang, et al. Illuminating protein space with a programmable generative model. Nature, 623:1070–1078, 2023. doi: 10.1038/s41586-023-06728-8.

[10] Brian L. Trippe, Jason Yim, Doug Tischer, David Baker, Tamara Broderick, Regina Barzilay, and Tommi Jaakkola. Diffusion probabilistic modeling of protein backbones in 3D for the motif-scaffolding problem, 2022. URL https://arxiv.org/abs/2206.04119. International Conference on Learning Representations (ICLR) 2023.

[11] Kieran Didi, Francisco Vargas, Simon V. Mathis, Vincent Dutordoir, Emile Mathieu, Urszula J. Komorowska, and Pietro Liò. A framework for conditional diffusion modelling with applications in motif scaffolding for protein design, 2024. URL https://arxiv.org/abs/2312.09236.

[12] Bram Mylemans, Boguslawa Korona, Amanda M. Acevedo-Jake, Ailsa MacRae, Thomas A. Edwards, Danny T. Huang, Andrew J. Wilson, Laura S. Itzhaki, and Derek N. Woolfson. De novo-designed bifunctional proteins for targeted protein degradation. Journal of the American Chemical Society, 148(32):34311–34319, 2026. doi: 10.1021/jacs.6c07593.

[13] Sandra Alonso, Gerard Santiago, Isabel Cea-Rama, Laura Fernandez-Lopez, Cristina Coscolin, Jan Modregger, Anna K. Ressmann, Mónica Martínez-Martínez, Helena Marrero, Rafael Bargiela, Marcos Pita, Jose L. Gonzalez-Alfonso, Manon L. Briand, David Rojo, Coral Barbas, Francisco J. Plou, Peter N. Golyshin, Patrick Shahgaldian, Julia Sanz-Aparicio, Víctor Guallar, and Manuel Ferrer. Genetically engineered proteins with two active sites for enhanced biocatalysis and synergistic chemo- and biocatalysis. Nature Catalysis, 3(3):319–328, 2020. doi: 10.1038/s41929-019-0394-4.

[14] Sergi Roda, Laura Fernandez-Lopez, Marius Benedens, Alexander Bollinger, Stephan Thies, Julia Schumacher, Cristina Coscolín, Masoud Kazemi, Gerard Santiago, Christoph G. W. Gertzen, Jose L. Gonzalez-Alfonso, Francisco J. Plou, Karl-Erich Jaeger, Sander H. J. Smits, Manuel Ferrer, and Víctor Guallar. A plurizyme with transaminase and hydrolase activity catalyzes cascade reactions. Angewandte Chemie International Edition, 61(37):e202207344, 2022. doi: 10.1002/anie.202207344.

[15] Karla M. Castro, Joseph L. Watson, Jue Wang, Joshua Southern, Reyhaneh Ayardulabi, Sandrine Georgeon, Stéphane Rosset, David Baker, and Bruno E. Correia. Accurate single-domain scaffolding of three nonoverlapping protein epitopes using deep learning. Nature Chemical Biology, 22(4):604–611, 2025. doi: 10.1038/s41589-025-02083-z.

[16] Xiaoran Jiao, Weian Mao, Zheng Sun, Yu Bo, Peiyuan Yang, Hao Chen, and Chunhua Shen. MoDiff: A protein diffusion model for multi-motif scaffolding problem. OpenReview preprint, 2024. URL https://openreview.net/forum?id=jXy5B0auu0. Posted October 4, 2024.

[17] Ke Liu, Weian Mao, Shuaike Shen, Xiaoran Jiao, Zheng Sun, Hao Chen, and Chunhua Shen. Floating anchor diffusion model for multi-motif scaffolding. In Ruslan Salakhutdinov, Zico Kolter, Katherine Heller, Adrian Weller, Nuria Oliver, Jonathan Scarlett, and Felix Berkenkamp, editors, Proceedings of the 41st International Conference on Machine Learning, volume 235 of Proceedings of Machine Learning Research, pages 31691–31708. PMLR, 2024. URL https://proceedings.mlr.press/v235/liu24av.html.

[18] Woody Ahern, Jason Yim, Doug Tischer, Saman Salike, Seth M. Woodbury, Donghyo Kim, In-drek Kalvet, Yakov Kipnis, Brian Coventry, Han Raut Altae-Tran, Magnus Bauer, Regina Barzilay, Tommi S. Jaakkola, Rohith Krishna, and David Baker. Atom-level enzyme active site scaffolding using RFdiffusion2. Nature Methods, 23:96–105, 2026. doi: 10.1038/s41592-025-02975-x. URL https://www.nature.com/articles/s41592-025-02975-x.

[19] Alex J. Li and Tanja Kortemme. All-atom protein design via SE(3) flow matching with ProteinZen. bioRxiv preprint, 2025. URL https://www.biorxiv.org/content/10.1101/2025.10.18.683228v1. Preprint posted October 18, 2025.

[20] Tomas Geffner, Kieran Didi, Zhonglin Cao, Danny Reidenbach, Zuobai Zhang, Christian Dallago, Emine Kucukbenli, Karsten Kreis, and Arash Vahdat. La-Proteina: Atomistic protein generation via partially latent flow matching, 2025. URL https://arxiv.org/abs/2507.09466.

[21] Jasper Butcher, Rohith Krishna, Raktim Mitra, Rafael I. Brent, Yanjing Li, Nathaniel Corley, Paul Kim, Jonathan Funk, Simon Mathis, Saman Salike, Aiko Muraishi, Helen Eisenach, Tuscan Rock Thompson, Jie Chen, Yuliya Politanska, Enisha Sehgal, Brian Coventry, Odin Zhang, Bo Qiang, Kieran Didi, Max Kazman, Frank DiMaio, and David Baker. De novo design of all-atom biomolecular interactions with RFdiffusion3. bioRxiv preprint, 2025. URL https://www.biorxiv.org/content/10.1101/2025.09.18.676967v1. Preprint posted September 18, 2025.

[22] Alexander E. Chu, Jinho Kim, Lucy Cheng, Gina El Nesr, Minkai Xu, Richard W. Shuai, and Po-Ssu Huang. An all-atom protein generative model. Proceedings of the National Academy of Sciences, 121(27):e2311500121, 2024. doi: 10.1073/pnas.2311500121.

[23] Zhuoqi Zheng, Bo Zhang, Kieran Didi, Kevin K. Yang, Jason Yim, Joseph L. Watson, Hai-Feng Chen, and Brian L. Trippe. MotifBench: A standardized protein design benchmark for motif-scaffolding problems, 2025. URL https://arxiv.org/abs/2502.12479.

[24] Justas Dauparas, Ivan Anishchenko, Nathaniel Bennett, Hua Bai, Robert J. Ragotte, Lukas F. Milles, Basile I. M. Wicky, Alexis Courbet, Robert J. de Haas, Neville Bethel, Paul J. Y. Leung, Timothy F. Huddy, Sam Pellock, Doug Tischer, F. Chan, B. Koepnick, H. Nguyen, Kang, B. Sankaran, A. K. Bera, N. P. King, and David Baker. Robust deep learning– based protein sequence design using ProteinMPNN. Science, 378(6615):49–56, 2022. doi: 10.1126/science.add2187.

[25] Michel van Kempen, Stephanie S. Kim, Charlotte Tumescheit, Milot Mirdita, Jeongjae Lee, Cameron L. M. Gilchrist, Johannes Söding, and Martin Steinegger. Fast and accurate protein structure search with Foldseek. Nature Biotechnology, 42:261–271, 2024. doi: 10.1038/s41587-023-01773-0.

[26] J. B. Kruskal. Multidimensional scaling by optimizing goodness of fit to a nonmetric hypothesis. Psychometrika, 29(1):1–27, 1964. doi: 10.1007/BF02289565.

[27] Yaron Lipman, Ricky T. Q. Chen, Heli Ben-Hamu, Maximilian Nickel, and Matt Le. Flow matching for generative modeling, 2023. URL https://arxiv.org/abs/2210.02747. International Conference on Learning Representations (ICLR) 2023.

[28] Jason Yim, Brian L. Trippe, Valentin De Bortoli, Emile Mathieu, Arnaud Doucet, Regina Barzilay, and Tommi Jaakkola. SE(3) diffusion model with application to protein backbone generation, 2023. URL https://arxiv.org/abs/2302.02277. International Conference on Machine Learning (ICML) 2023.

[29] Jason Yim, Andrew Campbell, Andrew Y. K. Foong, Michael Gastegger, José Jiménez-Luna, Sarah Lewis, Victor Garcia Satorras, Bastiaan S. Veeling, Regina Barzilay, Tommi Jaakkola, and Frank Noé. Fast protein backbone generation with SE(3) flow matching, 2023. URL https://arxiv.org/abs/2310.05297. Preprint under review; version 2 posted October 10, 2023.

[30] Avishek Joey Bose, Tara Akhound-Sadegh, Guillaume Huguet, Kilian Fatras, Jarrid Rector-Brooks, Cheng-Hao Liu, Andrei Cristian Nica, Maksym Korablyov, Michael Bronstein, and Alexander Tong. SE(3)-stochastic flow matching for protein backbone generation, 2024. URL https://arxiv.org/abs/2310.02391. International Conference on Learning Representations (ICLR) 2024.

[31] Jason Yim, Andrew Campbell, Emile Mathieu, Andrew Y. K. Foong, Michael Gastegger, José Jiménez-Luna, Sarah Lewis, Victor Garcia Satorras, Bastiaan S. Veeling, Frank Noé, Regina Barzilay, and Tommi S. Jaakkola. Improved motif-scaffolding with SE(3) flow matching, 2024. URL https://arxiv.org/abs/2401.04082. Transactions on Machine Learning Research (TMLR) 2024.

[32] Andrew Campbell, Jason Yim, Regina Barzilay, Tom Rainforth, and Tommi Jaakkola. Generative flows on discrete state-spaces: enabling multimodal flows with applications to protein codesign, 2024. URL https://arxiv.org/abs/2402.04997. International Conference on Machine Learning (ICML) 2024.

[33] Shiyu Zhu, Snigdha Gulati, Yang Liu, Sanket Kotnis, Qian Sun, and Yang Shen. Multimodal diffusion for joint design of protein sequence and structure. Protein Science, 34(12):e70340, 2025. doi: 10.1002/pro.70340.

[34] John Jumper, Richard Evans, Alexander Pritzel, Tim Green, Michael Figurnov, Olaf Ronneberger, et al. Highly accurate protein structure prediction with AlphaFold. Nature, 596: 583–589, 2021. doi: 10.1038/s41586-021-03819-2.

[35] Zeming Lin, Halil Akin, Roshan Rao, Brian Hie, Zhongkai Zhu, Wenting Lu, Nikita Smetanin, Robert Verkuil, Ori Kabeli, Yaniv Shmueli, Allan dos Santos Costa, Maryam Fazel-Zarandi, Tom Sercu, Salvatore Candido, and Alexander Rives. Evolutionary-scale prediction of atomic-level protein structure with a language model. Science, 379(6637):1123–1130, 2023. doi: 10.1126/science.ade2574.

[36] Yang Zhang and Jeffrey Skolnick. Scoring function for automated assessment of protein structure template quality. Proteins: Structure, Function, and Bioinformatics, 57(4):702–710, 2004. doi: 10.1002/prot.20264.

[37] Mihaly Varadi, Stephen Anyango, Mandar Deshpande, Sreenath Nair, Cindy Natassia, Galabina Yordanova, et al. AlphaFold protein structure database: massively expanding the structural coverage of protein-sequence space with high-accuracy models. Nucleic Acids Research, 50(D1): D439–D444, 2022. doi: 10.1093/nar/gkab1061.

